# The effective family size of immigrant founders predicts their long-term demographic outcome: from Québec settlers to their 20th-century descendants

**DOI:** 10.1101/2021.07.25.453708

**Authors:** Damian Labuda, Tommy Harding, Emmanuel Milot, Hélène Vézina

## Abstract

Human evolution involves population splits, size fluctuations, founder effects, and admixture. Population history reconstruction based on genetic diversity data routinely relies on simple demographic models while projecting the past. No specific demographic assumptions are needed to understand the genetic structure of the founder population of Québec. Because genealogy and genetics are intimately related, we used descending genealogies of this population to pursue the fate of its founder lineages. Maternal and paternal lines reflect the transmission of mtDNA and the Y-chromosome, respectively. We followed their transmission in real-time, from the 17^th^ century down to its 20^th^-century population. We counted the number of married children of immigrants (i.e., their effective family size, EFS), estimated the proportion of successful immigrants in terms of their survival ratio, and assessed net growth rates and extinction. Likewise, we evaluated the same parameters for their Québec-born descendants. The survival ratio of the first immigrants was the highest and declined over time in association with the decreasing immigrants’ EFS. Parents with high EFS left plentiful married progeny, putting EFS as the most important variable determining the parental demographic success throughout time for generations ahead. The 17^th^ and 18^th^-century immigrants bear the most remarkable demographic and genetic impact on the 20^th^-century population of Québec. Lessons learned from Québec genealogies can teach us about the consequences of founder effects and migrations through real people’s history. The effective family size of immigrant founders predicts their long-term demographic outcome.

## INTRODUCTION

Population genetics is about population structure, genetic history, and phylogenetic connections. It relies on likelihoods using demographic models that assume long-term equilibria. To reveal populations’ genetic past and demographic history, we routinely model populations’ ascending genealogical trees (1). However, “the history of the human species and of particular populations suggests that long-term equilibria, an assumption of many genetic models, is only a convenient mathematical fiction” (2, 3) (and see also (4)). In this context, genealogies represent an objective data source to learn about populations’ demographic past and their ensuing genetic structure. Their value is favorably enhanced once merged with current genetic diversity data (5-9). Therefore, studies of populations with an extensive genealogical record may fill the gap between historical reality and computer modeling. However, genealogies do not exceed historical depth but usually represent firm data. In contrast, those projected using genetic diversity data (phylogenetic genealogies or *gene trees*) are scaled by a mutational clock and the effective population size and head back to prehistory. Importantly, lessons are to be learned about mechanisms driving populations’ evolution at prehistorical scales from the generations’ scale data using genealogical records.

Genealogical coverage exists only for a handful of human populations, such as small isolates or local scale data on religious groups (10-13) and Island (5). The Québec population is presumably the largest one among these genealogically-covered populations. European settlement started in the St-Lawrence Valley in the 17^th^ century, primarily by immigrants from France (14, 15). BALSAC is a genealogical database of the population of Québec, which includes data on married individuals from the start of the colony up to 1960. The descending genealogies comprise information on the number of immigrants and their immediate descendants, their demographic performance, sex ratio, and other essential characteristics and how these changed with time. Notably, they also include records on married individuals that left no descendants. Thus, there are no direct genetic traces in the present-day diversity data that are crucial to understanding the population demographic past. Descending genealogies allow studies on evolutionary and selection events (16, 17), geographical mapping of disease-causing mutations (18, 19), and many more.

Genealogical lines relating mother-and-daughter or father-and-son reflect the transmission of the uniparentally inherited mitochondrial DNA and non-recombining portion of the Y-chromosome, respectively. There is no ambiguity in their reconstruction, reflecting a lack, at the genetic level, of gametic sampling and recombination. Thus, we can separately analyze the effects of sex-asymmetry in reproductive success, immigration, and generation time. And indeed, the first genetic studies to learn about populations past exploited DNA diversity by following uniparentally inherited mtDNA and the Y-chromosome. Rarely genetic studies focused on maternal mtDNA and paternal Y-chromosome diversity at the same time (20). Their joint analyses relying on their genetic diversities are complicated because the mutation rates of mtDNA and the Y-chromosome are very different (21). These analyses are further complicated by differences in the effective population size of males and females. This problem disappears once we follow maternal and paternal genealogical lines and their distribution among generations down the road (5). Therefore, we can concurrently analyze maternal and paternal lines using genealogical data, disregarding any mutational bias.

In this study, we use the descending genealogy of married individuals to follow the fate of maternal and paternal lineages over a period covering more than three hundred years. Our goal is to assess how historical immigration waves shaped the genetic structure of the Québec population and to understand the contemporary genetic legacy of Quebec lineage founders as a function of their immigration history and the reproductive success of their descendants. We describe the overall dynamics of different immigration waves relying on new statistics that include information from descending genealogies, unavailable in earlier population history studies. Using this information, we separately analyze the effects of sex-asymmetry in reproductive success, immigration, and generation time.

## MATERIALS AND METHODS

### BALSAC data

Genealogical information on the historical Québec population is digitalized in the BALSAC population database (http://balsac.uqac.ca) with a remarkably high degree of completeness. BALSAC includes data from the 17^th^ and 18^th^ centuries in Québec that were initially compiled within the Early Québec Population Register of the Programme de recherche en démographie historique (PRDH) of the Université de Montréal (22) (https://www.prdh-igd.com/en/home). The genealogies were primarily reconstructed from Catholic church records with information about life-history events such as marriages, births, or deaths. Moreover, BALSAC and PRDH collected valuable information on genealogical connections beyond Québec, such as data on European and Acadian ancestors or emigrant marriages outside Québec (e.g. (Tremblay and Vezina 2010, Moreau, Vezina et al. 2011)). We define immigrants as the first generation of settlers to marry in Quebec; therefore, some were born in Quebec to parents who married before coming to Québec. Most immigrants introduced new maternal and paternal lineages, representing new mitochondrial DNA and Y-chromosome lines, respectively, and are recognized as lineage founders. However, some immigrants were related to each other upon their arrival in Québec (e.g., siblings, cousins). In those cases, the most recent common ancestor (maternal or paternal) of these related immigrants was identified as the lineage founder to avoid counting a lineage more than once (Figure S1 in Supplementary Material). The BALSAC register is almost complete for individuals married in the Catholic parishes before the 1960s and partial for other confessions (23). Therefore, the population of married couples between 1931 and 1960 was set as the reference for the 20^th^-century population of Québec. This 30-year period is approximately the time of one generation (24).

### Demographic parameters

From the BALSAC database, we extracted all married individuals and categorized them either as immigrants or as Québec-born (QB) descendants of the immigrants (c.f. Figure S1). Also, unless stated otherwise, immigrant males and females were counted independently to separate maternal and paternal lineages and pooled into waves *w*_*i*_ corresponding to their marriage years *t*_*i*_, starting at 1621 and up to 1930. QB were grouped into corresponding strata *s*_*i*_, according to their marriage year *t*_*i*_.

*N*_*i*_ represents the count of immigrants from wave *w*_*i*_ in the year *t*_*i*_ (both sexes, or only either males or females, according to the context). *Nw*_*i*_ denotes the number of their male or female descendants married between 1931 and 1960 (note that only maternal lineages in married women are counted: male mtDNA carriers are excluded because they do not transmit mtDNA further). Their ancestors are *contributing* individuals, namely, immigrants or Québec-born, whose lineage survived until 1931-60, respectively designated by *Nc*_*i*_ and *Nc*_QB*i*_. Likewise, *N*_QB*i*_ represents the count of *s*_*i*_ QB married in year *t*_*i*_ and *Ns*_QB*i*_ that of their descendants married in Québec between 1931-1960.

To assess the overall rate of the demographic growth *k* in Québec, we used historical estimates of the Québec population size from Larin (25), census reports (26), and BALSAC data (only considering married individuals). With a growing population, *N*_*t*_=*N*_*t-x*_ (1+*k*)^*x*^, with *N*_*t*_ being the population size at the year *t, N*_*t-x*_, its size at *x* years earlier, and *k* is the population growth rate per year. Because *k* is small, (1+*k*) ^*x*^ *∼ e* ^*k* x^, it follows that ln(*N*_*t*_)-ln(*N*_*t-x*_)=*k·x*, and so the population growth rate *k* can be readily estimated from the slope of ln *N*_*t*_ versus time *t* (Suppl Figure S2).

### New descriptors of descending genealogies

Remarkably, descending genealogies come with additional information that is not captured in the ascending genealogies. It includes the number of initial founders, that of their descendants and portion of the surviving (and lost lineages). We introduce three measures to describe this information on population demographic history, namely: (i) the survival ratio (SR_*i*_), (ii) the net growth (*dc*_*i*_), and (iii) net extinction (*a*_*i*_) rates. SR_*i*_ = *Nc*_*i*_ /*N*_*i*_ for immigrants and *Nc*_QB*i*_/*N*_QB*i*_ for Québec-born individuals, thereby describing the proportion of contributing individuals within *w*_*i*_ or *s*_*i*_. The net growth rate *dci* and extinction rate *ai*, between historical *t*_*i*_, and the descendant population of the 1931-60 Québec is estimated using *t* =1947.5 (*t*_*1947*.*5*_) as an average target year of marriages between1931-60. Both *dci* and *ai* are evaluated using the same rate equation we used to estimate *k*, except that they only describe a net growth or extinction of the population fraction from time *t*_*i*_ down to the average *t*=1947.5. Thus, we write *Nw*_*i*_=*Nc*_*i*_(1+*dc*_*i*_)^(*t*^_*1947*.*5*_^*-ti*)^, whereby *dc*_*i*_ = ln(*Nw*_*i*,_/*Nc*_*i*_)/*(t*_*1947*.*5*_*-t*_*i*_) and, likewise, *dc*_QB*i*_ = ln(*Ns*_QB*i*_ /*Nc*_QB*i*_)/(*t*_*1947*.*5*_-*t*_*i*_). As for the net extinction rate for immigrants: *Nc*_*i*_=*N*_*i*_(1-*a*_*i*_)^*(t*^_*1947*.*5*_^*-ti*)^ and *a*_*i*,_ = ln(*N*_*i*_/*Nc*_*i*_)/(*t*_*1947*.*5*_-*t*_*i*_) and likewise for the Québec-born *a*_QB*i*,_ = ln(N_QB*i*_/*Nc*_QB*i*_)/(*t*_*1947*.*5*_*-t*_*i*_). Note that *dc*_*i*_ and *a*_*i*_ complement SR_*i*_ and 1-SR_*i*_, respectively, but also consider the time factor (*t*-*t*_*i*_). To plot the figures, we pooled data in bins of five or ten years to reduce the variance, yet the values shown are averages per year.

### Progeniture

We refer to the number of married children of a couple as their effective family size (EFS) (27) or the number of potentially fertile children (i.e., grand-child bearing children) (5, 16, 27). While BALSAC data does not allow to calculate Darwinian fitness of individuals directly, fortunately, the EFS is strongly correlated to fitness in the French-Canadian population and was used here as a proxy (16, 27). However, (28) see the danger of conflating selection and inheritance with such a fitness proxy, although the strong correlation limits this problem. We measured EFS sex-wise by counting the number of married daughters (EFS-daughters) or sons (EFS-sons) for mothers and fathers, respectively, or by pooling both sexes’ maternal and paternal children. We separately examined the EFS of contributing and non-contributing parents from immigrants or their Québec-born descendants.

## RESULTS

### Population founders and their descendants in 1931-60 Québec

The European founding of the Québec population started with the establishment of Québec City in 1608, followed by urban centers at Trois-Rivières (1634) and Montreal (1642), upstream of the Saint Lawrence River. The settlements in Beaupré (1636) and Baie-Saint-Paul (1673), north-east of Québec City, largely contributed to the peopling further downstream. In 1763, the Treaty of Paris sealed the British conquest of Québec and Canada. Around 1800, the Québec population had grown to about 220,000 people (25), eventually reaching 2,874,662 in 1931 (26). In BALSAC (version as of March 2015), we find genealogies of 3,340,072 individuals married in Québec before 1961 (2,018,038 before 1931). This includes 455,687 immigrants, of whom 276,946 settled before 1931. Of the latter, 29,668 immigrant women and 32,608 immigrant men contributed to the uniparentally-inherited lines of 1931-60 Québec (Table 1). In its early years, the Québec population primarily grew due to immigrant settlers (Figure 1A; Table 1; Figure S2). The first immigrant marriage documented in Québec Catholic registers took place in 1621.

**Table 1.** Numbers of immigrants and Québec-born individuals who married in Quebec in five historical periods. Numbers of distinct and novel (newly introduced, i.e., in addition to those already present and introduced earlier), uniparentally inherited lineages (separately maternal and paternal) are reported as well. Contributing immigrants are those who transmitted their descendant lineages in the 1931-60 population.

| Periods | 1620-1680 |  | 1681-1750 |  | 1751-1800 |  | 1801-1850 |  | 1851-1930 |  | Sum down to 1930 |  | 1931-60 |  |
| --- | --- | --- | --- | --- | --- | --- | --- | --- | --- | --- | --- | --- | --- | --- |
|  | Women | Men | Women | Men | Women | Men | Women | Men | Women | Men | Women | Men | Women | Men |
| **Immigrants** | 1241 | 1631 | 1580 | 4827 | 3981 | 6766 | 26749 | 27515 | 99518 | 103138 | 133069 | 143877 | 81565 | 97176 |
| Distinct lineages | 1081 | 1520 | 988 | 4101 | 2567 | 5345 | 22781 | 24154 | 75553 | 86899 | 102094 | 121221 | 61524 | 79543 |
| Novel lineages | 1081 | 1520 | 930 | 4052 | 2543 | 5279 | 22696 | 24030 | 74844 | 86340 | 102094 | 121221 | 57961 | 76357 |
| **Contributing immigrants** | 706 | 827 | 470 | 1753 | 922 | 1816 | 2117 | 2344 | 25453 | 25868 | 29668 | 32608 | 81565 | 97176 |
| Distinct lineages | 616 | 793 | 238 | 1440 | 331 | 1281 | 1914 | 2106 | 20923 | 22777 | 23873 | 28233 | 61524 | 79543 |
| Novel lineages | 616 | 793 | 206 | 1416 | 321 | 1255 | 1878 | 2046 | 20852 | 22723 | 23873 | 28233 | 57961 | 76357 |
| **Quebec-born** | 312 | 56 | 14726 | 11135 | 43526 | 37891 | 146715 | 134197 | 707013 | 645521 | 912292 | 828800 | 586049 | 557244 |
| **Contributing Quebec-born** | 286 | 49 | 13987 | 10272 | 42871 | 36197 | 144000 | 130520 | 689709 | 630727 | 890853 | 807765 | 580335 | 553516 |
| **Immigrants + Quebec-born** | 1553 | 1687 | 16306 | 15962 | 47507 | 44657 | 173464 | 161712 | 806531 | 748659 | 1045361 | 972677 | 667614 | 654420 |
| **Fraction of immigrants** | 0.80 | 0.97 | 0.10 | 0.30 | 0.08 | 0.15 | 0.15 | 0.17 | 0.12 | 0.14 | 0.13 | 0.15 | 0.12 | 0.15 |

**Figure 1.**
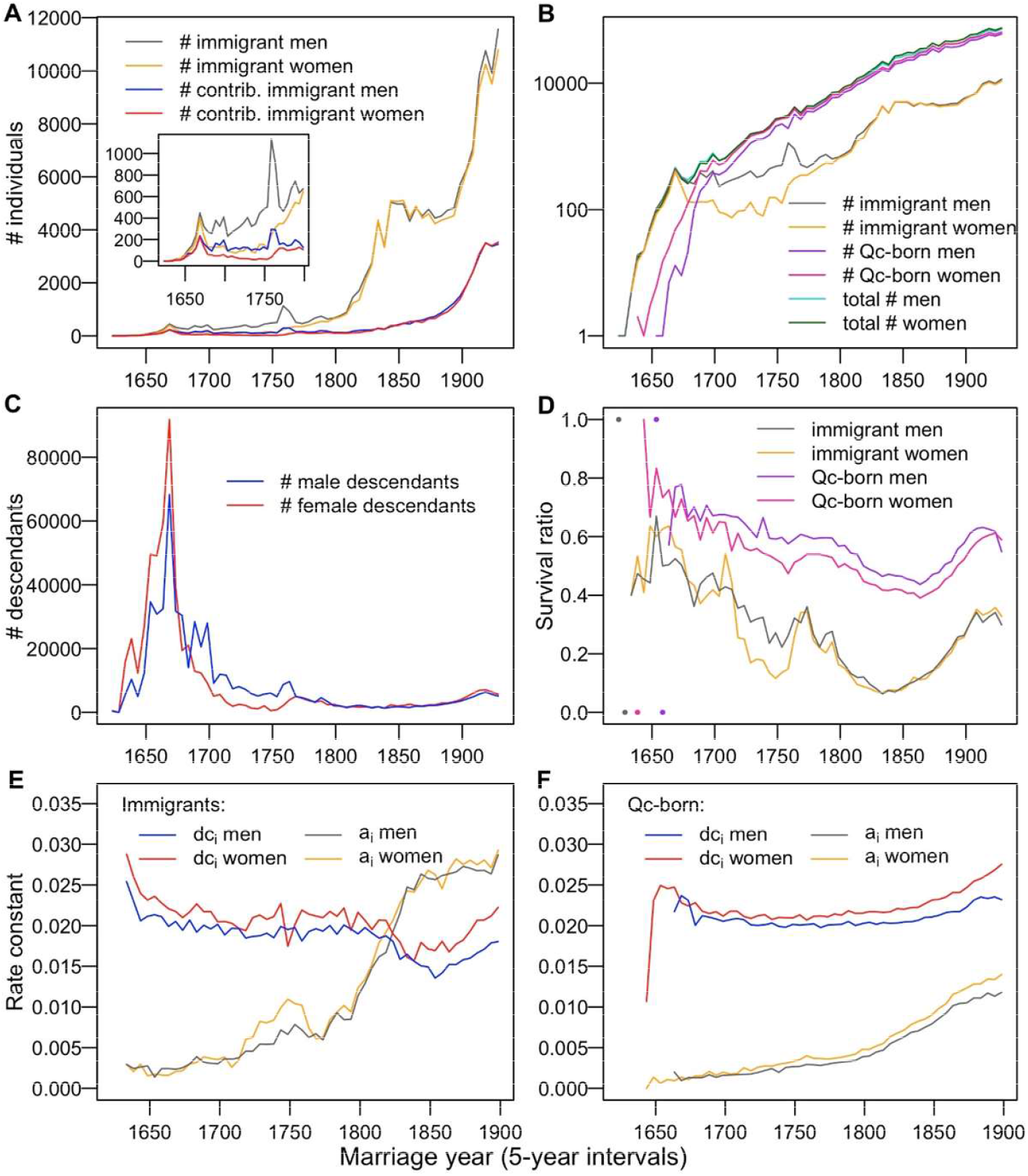
Immigrants, their Quebec-born descendants and related statistics as a function of time. The number of all immigrants and their contributing portion at linear scale (inset shows early history at a magnified scale) (**A**). The number of all immigrants, Québec-born and their total in logarithmic scale (**B)**. The number of immigrants’ descendants, *Nw*_*i*_, within the 1931-60 population as a function of immigrants’ settlement time (**C**). Evolution of the survival ratio SR of immigrants and Quebec-born (**D**). Corresponding plots of the net growth (*dc*_*i*_) and net extinction (*a*_*i*_) rates for the immigrants (**E**) and the Quebec-born (**F**).

### Historical periods and population expansion

Based on historical characteristics, we divided the European settling of the Quebec territory into five immigration periods. From 1621 to 1680, the first period was marked by the arrival of the Filles du Roy in the years 1663-73. They were French women whose immigration to New France was sponsored by King Louis XIV (thus “King’s daughters”), designed to boost New France’s population. They represent the first immigration peak in Figure 1A. About twelve hundred women and sixteen hundred men immigrants were registered at that period. Out of these, 708 women and 830 men left descendant lineages (Table 1) that account for 66.6% of maternal and 47.3% paternal lineages in the 1931-60 Québec population l (Table 2).

**Table 2.**
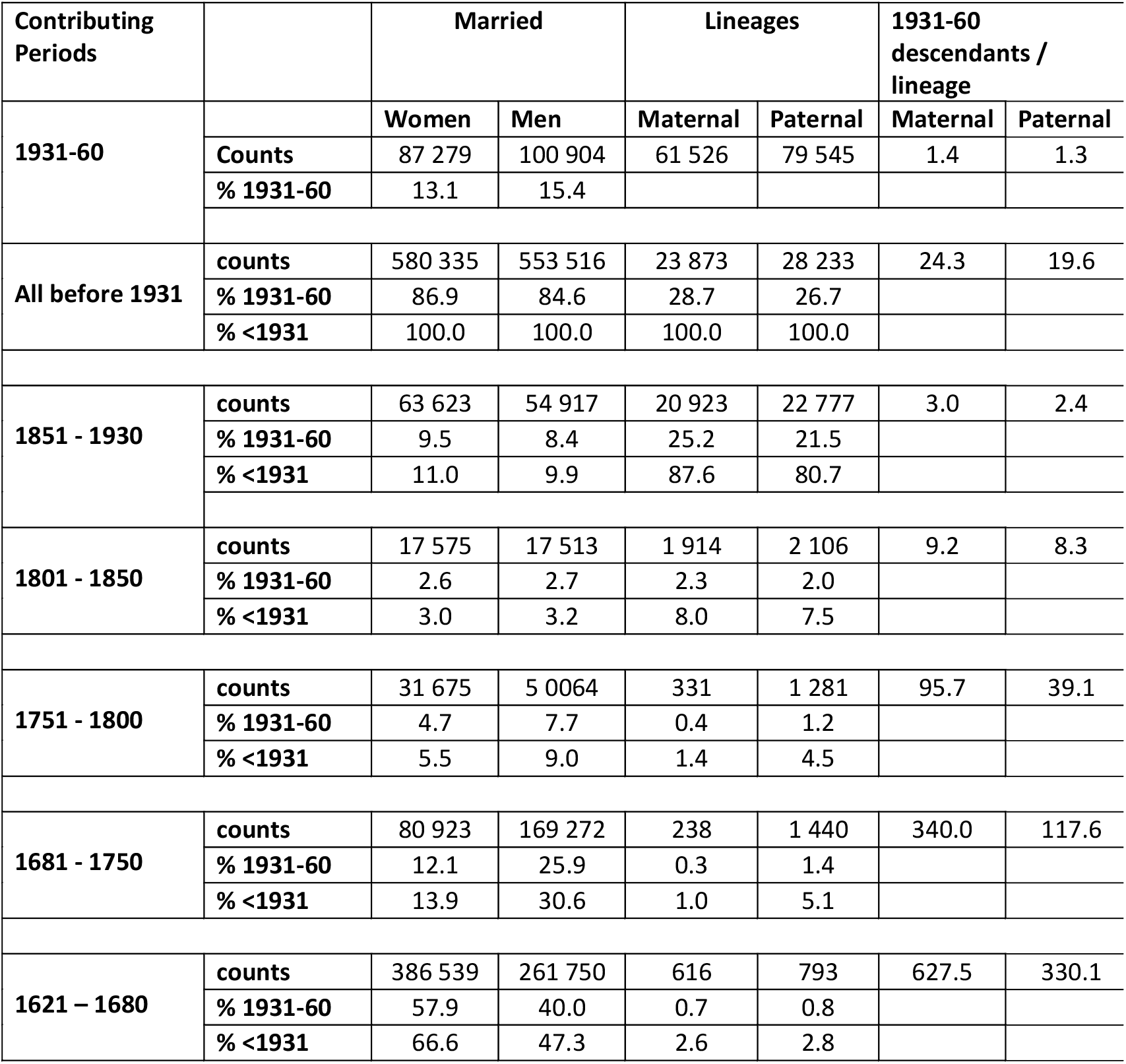
Contribution of immigrants from different periods to the 1931-60 Québec, both in the number of their married descendants and in the number of distinct maternal and paternal lineages.

| Contributing Periods |  | Married |  | Lineages |  | 1931-60 descendants / lineage |  |
| --- | --- | --- | --- | --- | --- | --- | --- |
|  |  | Women | Men | Maternal | Paternal | Maternal | Paternal |
| 1931-60 | Counts | 87 279 | 100 904 | 61 526 | 79 545 | 1.4 | 1.3 |
|  | % 1931-60 | 13.1 | 15.4 |  |  |  |  |
| All before 1931 | counts | 580 335 | 553 516 | 23 873 | 28 233 | 24.3 | 19.6 |
|  | % 1931-60 | 86.9 | 84.6 | 28.7 | 26.7 |  |  |
|  | % <1931 | 100.0 | 100.0 | 100.0 | 100.0 |  |  |
| 1851 - 1930 | counts | 63 623 | 54 917 | 20 923 | 22 777 | 3.0 | 2.4 |
|  | % 1931-60 | 9.5 | 8.4 | 25.2 | 21.5 |  |  |
|  | % <1931 | 11.0 | 9.9 | 87.6 | 80.7 |  |  |
| 1801 - 1850 | counts | 17 575 | 17 513 | 1 914 | 2 106 | 9.2 | 8.3 |
|  | % 1931-60 | 2.6 | 2.7 | 2.3 | 2.0 |  |  |
|  | % <1931 | 3.0 | 3.2 | 8.0 | 7.5 |  |  |
| 1751 - 1800 | counts | 31 675 | 5 0064 | 331 | 1 281 | 95.7 | 39.1 |
|  | % 1931-60 | 4.7 | 7.7 | 0.4 | 1.2 |  |  |
|  | % <1931 | 5.5 | 9.0 | 1.4 | 4.5 |  |  |
| 1681 - 1750 | counts | 80 923 | 169 272 | 238 | 1 440 | 340.0 | 117.6 |
|  | % 1931-60 | 12.1 | 25.9 | 0.3 | 1.4 |  |  |
|  | % <1931 | 13.9 | 30.6 | 1.0 | 5.1 |  |  |
| 1621 – 1680 | counts | 386 539 | 261 750 | 616 | 793 | 627.5 | 330.1 |
|  | % 1931-60 | 57.9 | 40.0 | 0.7 | 0.8 |  |  |
|  | % <1931 | 66.6 | 47.3 | 2.6 | 2.8 |  |  |

The second immigration cycle in Quebec, from 1681 to 1750, took place after the arrival of Filles du Roy and before the British conquest (1760). <u>T</u>his period contributed 13.9% and 30.6% of maternal and paternal descendants to the 1931-60 population. The paternal contribution prevailed over the maternal lineages. Altogether, the uniparentally inherited lines introduced by the immigrants before 1751 are found in ∼467,000 women and ∼431,000 men married between 1931 and 1960 (80.5 % and 77.9 % of all married descendants from before 1931, respectively; Table 2). Note that at the turn of the 18^th^ century, QB individuals surpassed the immigrants in numbers and started to dominate the bulk of the population (Figure 1B).

From 1751 to 1800, the third colonization period is marked by a remarkable peak of male immigrants (Table 2; Fig. 1A) due to the demobilization of French soldiers (25), who settled in Québec after the British takeover in 1760. They were joined by descendants of French pioneers from Acadia (present-day Nova Scotia and New Brunswick), who were survivors of the British deportation campaign of 1755. Acadian settlers were followed by British Loyalists fleeing the United States after the US Declaration of Independence. Immigration dramatically increased after 1800 (Figure 1A and B). It slowed down in the middle of the 19^th^ century and resumed at the turn of the 20^th^ century. About 55,000 immigrants arrived in Quebec during the 1801-1850 period (4th period) and more than 200,000 from 1851 to 1930 (5th period).

Up to 1680, the Québec population growth mainly reflects an increasing number of new settlers. Estimates of *k* for this period, based on Larin’s data (*k* = 0.084) and BALSAC data (*k* = 0.11), concord very well with each other, considering the relatively small number of individuals involved (Figure S2). During the 1681-1850 period, the population growth rate is estimated at 0.026 with both data sets (Larin and BALSAC) and *k*∼0.015 following 1850. The population growth after 1680 is primarily due to QB, while the contribution of the new immigrants dropped to about 15% (Figure 1B). In comparison, the average population growth rate in Europe between 1500 and 1900 was about 0.006 per year (29).

### Survival ratio, net growth, and extinction of lineages

While focusing on maternal and paternal lines, we counted how many lineages survived, how many were lost, how fast they grew, and what was their extinction rate; all this within the 1931-60 Québec population as a target. The demographic outcome of an immigration wave *i* among the 1931-60 descendants, *Nw*_*i*_ = *N*_*i*_[(1 − *a*_*i*_)(1 + *dc*_*i*_)]^(*t*1947−*ti*)^s a function of the immigrants ‘number *N*_*i*_, their arrival time *t*_*i*_, the extent of their growth (*dc*_*i*_), and lineage loss (*a*_*i*_).

The maximum *Nw*_*i*_ coincides with the immigration peak of the Filles du Roy (Figure 1C). Note that an excess of females *Nw*_*i*_ preceded and was within this maximum. This was followed by much greater male than female *Nw*_*i*_, reflecting the prevalence of male immigrants. In contrast to the 17^th^-18^th^ century immigrant contributions (Tables 1 and 2), the *Nw*_*i*_ of the 19^th^-century immigrants (Figure 1C) appears strikingly low for both sexes, despite an upsurge in immigrant numbers (Figure 1A and B) as judged by the relatively low level of descendants in 1931-60 Québec (Figure S3).

Before 1681, the survival ratio of immigrants was around 0.6 (Figure 1D). Its subsequent decrease can be associated with a drop in net growth, *dc*_*i*_, and an increase in net lineage extinction, *a*_*i*_. In the first half of the 19^th^ century, there is a sudden *dc*_*i*_ drop and a sharp rise in the immigrant *a*_*i*_. This combination explains the minimum immigrant survival ratio with a nadir of 0.1 at around 1835. Around 1825, *a*_*i*_ became larger than *dc*_*i*_, (i.e., ln(*N*_*i*_/*Nc*_*i*_) > ln(*Nw*_*i*_/*Nc*_*i*_), such that *N*_*i*_ > *Nw*_*i*_). It means that from that time on, the number of immigrants exceeded the number of their descendants among the 1931-60 marrieds (compare data in Tables 1 and 2).

### Effective family size - determinants of the outcome

The demographic outcome of immigrants depends on their reproductive performance and that of their descendant generations. The QB survival ratio follows a similar trajectory but with higher survival ratios and ranges from about 0.8 to a minimum of only 0.4 in the middle of the 19^th^ century (Figure 1D). The extinction rate *a*_*i*_ of QB paralleled that of the immigrants, albeit at a lesser pace. QB appear more demographically stable but still succumb to the same external factors, thereby decreasing the survival ratio and slowing down their growth. QB did not go through the 19^th^-century *dc*_*i*_ downfall that affected immigrants (Figure 1E and F). EFS values are the highest for married individuals in the 17^h^-century, followed by a minimum around 1750, a rebound at the turn of the 19^th^ century to a decline eventually. Contributing QB and immigrants show similar ups and downs in EFS (Figure 2). However, overall, QB have a greater EFS than the immigrants (Figure 2 – middle curves). Non-contributing individuals had a much lower EFS throughout the study period, with immigrants showing the lowest EFS with an average of ∼0.5, i.e., one married child per couple, either a boy or a girl (Figure 2).

**Figure 2.**
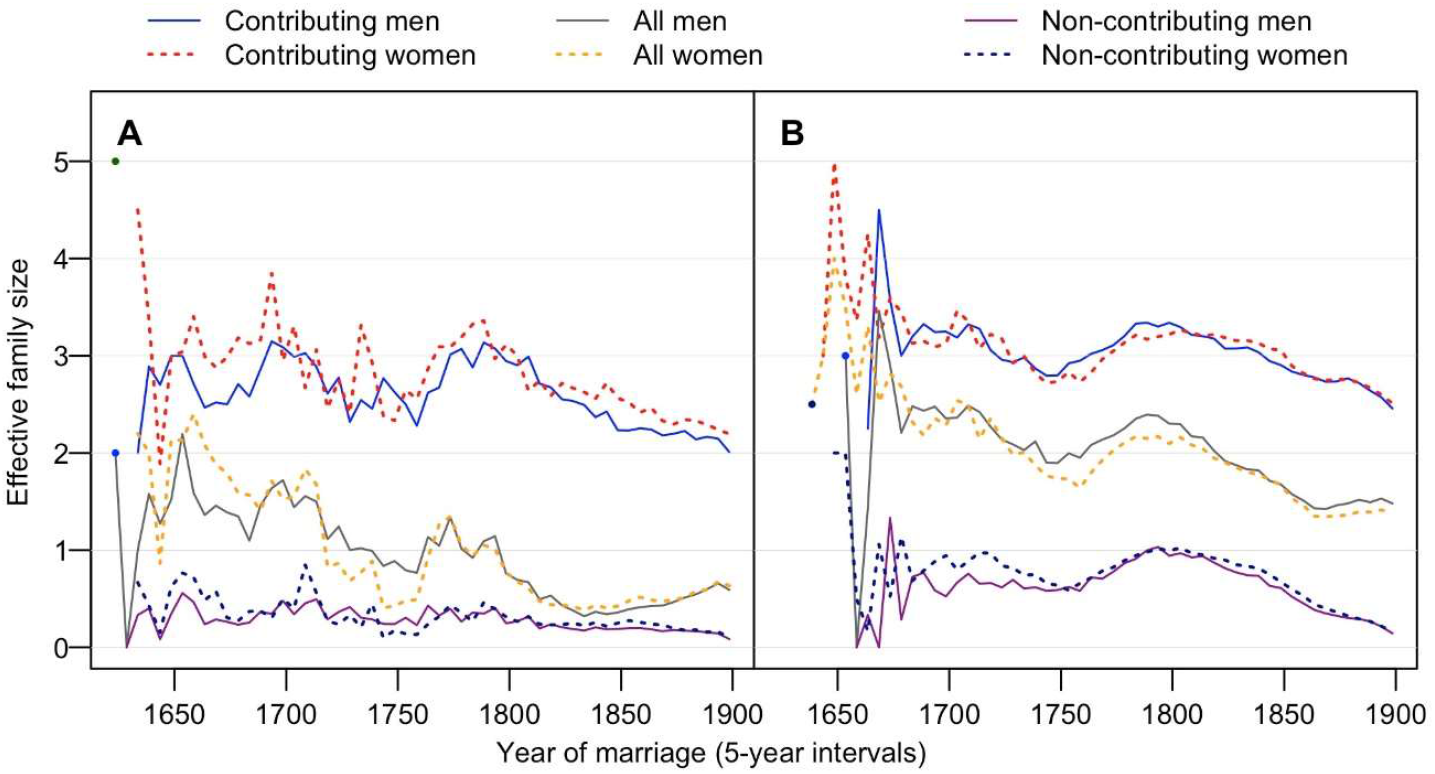
Effective family size of maternal and paternal lineages at different time points. Families of immigrants (**A**), and Quebec-born families (**B**), showing EFS of contributing, non-contributing lineages and their total (All) as indicated. Maternal EFSd are shown in dotted lines and paternal EFSs in solid lines.

Notably, the survival ratio of immigrants follows their EFS (Figures 3A and B). The survival ratio of the maternal and paternal lineages of the immigrants also strongly correlates with their corresponding EFS (R^2^ = 0.91 and 0.90, respectively; p << 0.001). In other words, the proportion of the contributing immigrants’ lineages from any time *i* mirrors their EFS. This correlation also stands when we consider 1-year rather than 5-year intervals, despite a larger variance (Figure S4). An immigrant with an EFS = 1 (1 child, either daughter or son) predicts that its descending lineage (maternal or paternal) has a ∼0.3 probability of surviving up to ≥1931 (insets of Fig. 3A and B). The correlation between the survival ratio and EFS is perpetuated over QB generations descending from these immigrants (Figure 3C and D; Figure S4), although the correlation is less straightforward (R^2^ = 0.65 and 0.69) (Figure 3C and D). And indeed, by the end of the 18^th^ century, QB EFS and SR curves start to diverge and then cross by the end of the 19^th^ century, thereby inverting the relation (see Discussion).

**Figure 3.**
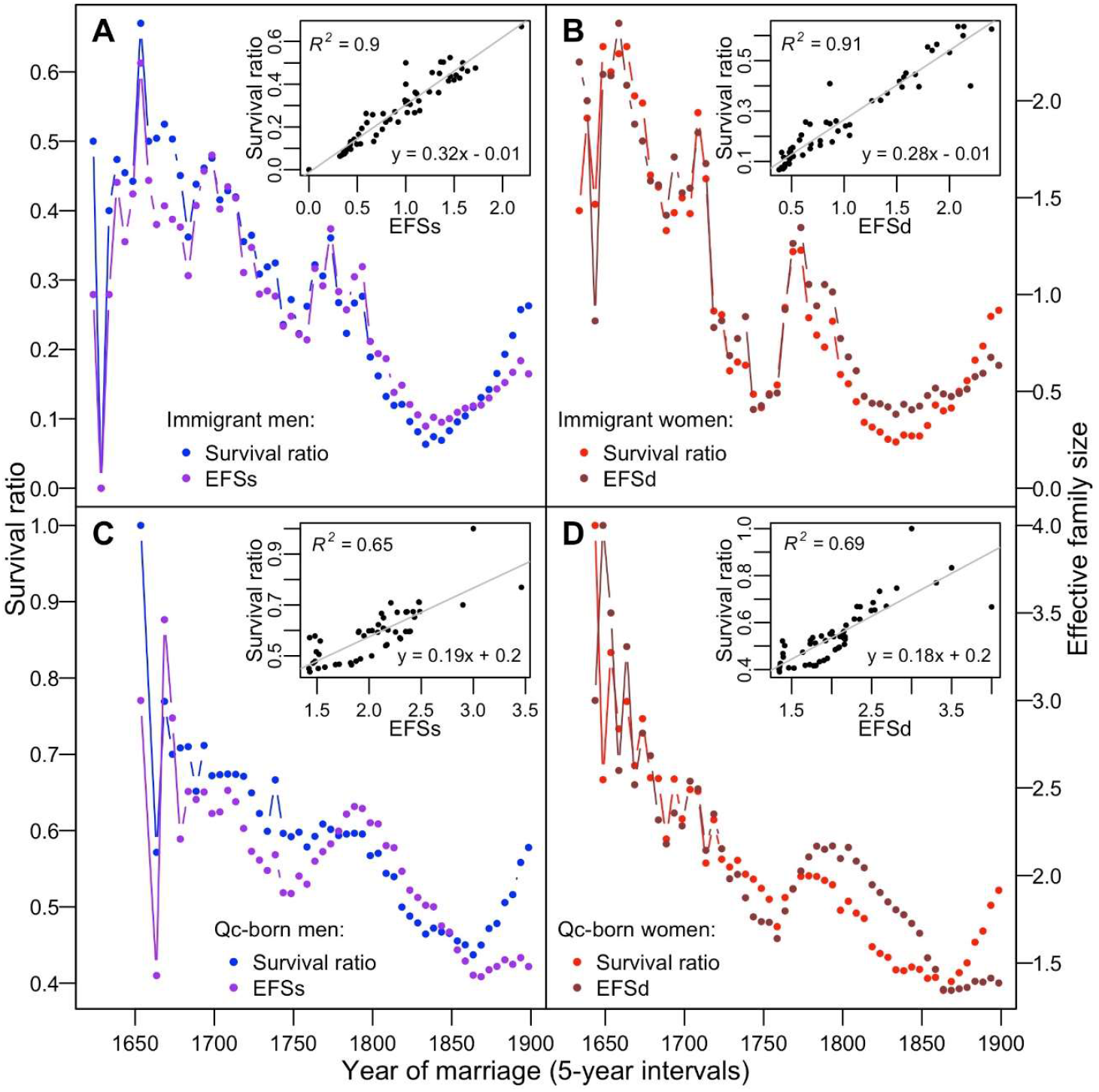
Correlation between effective family size and survival ratio. Overlapping plots of total EFSs and EFSd and their survival ratios for the immigrants (**A** and **B**) and the Quebec-born lineages (**C** and **D**). Their correlation plots are shown in the insets.

### Sex-asymmetry among immigrants

During the French Regime, there were fewer immigrant women than immigrant men. Before 1681, the immigrants’ sex ratio (men to women) was 1.3 (Table 1). It went up to higher values in the middle of the 18^th^-century (Figure S5) with an average of 2.1 between 1681 and 1800. At the beginning of the colony, women married much younger than men and remarried more often (15, 30). In Figure 4, we show that before 1681 remarriages of women prevailed and that this tendency reverted after 1681. During the following years, given the shortage of immigrant women, immigrant men married Québec-born daughters of previous immigrant couples. This created a disequilibrium between EFSd and EFSs of QB parents. According to BALSAC with the PRDH data included (see M&M), during the early time of Quebec (parental marriages before 1700), Québec-born males were less nuptially successful than their sisters (Table 1), as shown by the parents’ EFSs that was lower than EFSd (Figure 2 and S6). It is plausible that many QB males were lost from the “radar” of Québec church records; as explorers of new territories themselves, they became emigrants.

**Figure 4.**
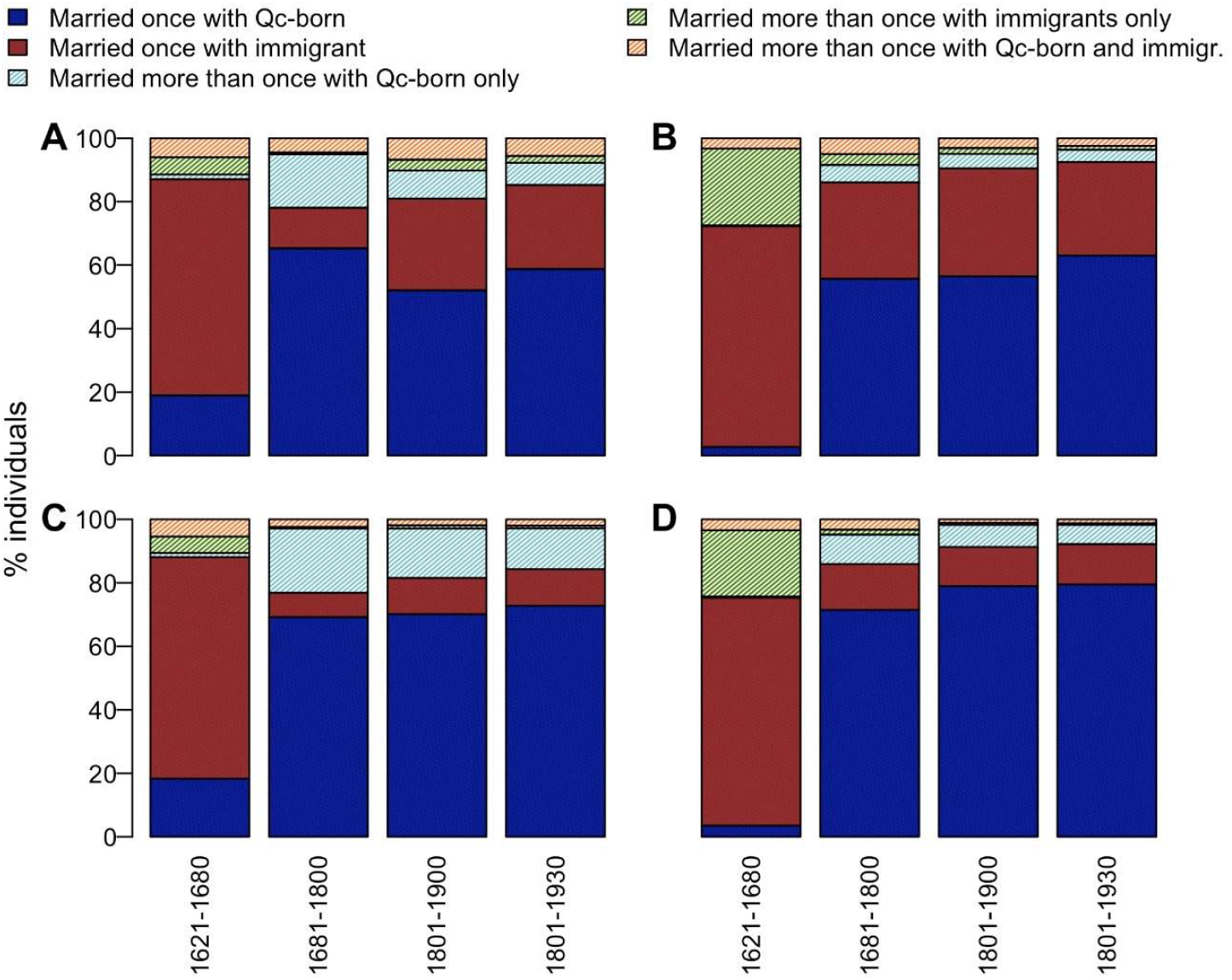
Distribution of marriages and remarriages among marrieds at four time periods. Different categories of marrieds are specified in the above histograms by their corresponding color codes. The contributing male and female immigrants are in **A** and **B**, respectively, whereas all (immigrants and Quebec-born) married men (**C**) and women (**D**).

Consequently, we observe the preferential impact of the matrilineal lineages from before 1681 (Figure 1C) because successive immigrant males marrying QB daughters promoted the expansion of maternal lines introduced earlier. (Note that years on the x-axis are identifying parental marriages, yet the statistics of the reporting children marriage, such as EFS, are taking place one generation later). At the same time, these male immigrants enriched the paternal lineages’ genetic pool relative to that of the ill-represented immigrant maternal lines (Figure 1C, Figure S7). In the 1931-60 population, we find 630 individuals per single maternal lineage introduced before 1681, compared to only 330 per single patrilineal line (Table 2). Considering all descendants from before 1801, we find three times more paternal than maternal lineages in the 1931-60 population, 3514 and 1185, respectively.

In contrast, over the 19^th^ century, down to 1930, there were 24 883 paternal and 22 837 maternal contributing immigrants (Table 2). The sex ratio of 1.09 concords with an excess of male remarriages compared to the period before 1681 when female remarriages prevailed (ratio of 0.92). Male immigration bias profoundly affected the frequency spectra of male and female lineages introduced before 1801 (Figure S7). At the molecular level lower female lines diversity was observed by contrasting the Y chromosome versus mtDNA diversities in Gaspésie (31) and that is well known from the demographic comparisons of the frequencies of the patronyms with that of the matronyms using genealogical information (32)

## Discussion

Turning to the coalescent (33, 34), we can use the existing genetic variation to reconstruct ascending genealogical and phylogenetic trees modeling the past based on today’s outcome. Having access to population-wide descending genealogies, we do not have to depend on demographic models and infer history from the present-day diversity (1). The BALSAC database, which consists of digitized vital records from Quebec for over three centuries, enables us to investigate the demographic past of the Québec population directly. Knowing the underlying history, we better understand the present. This paper analyzes uniparentally inherited maternal and paternal lineages of Quebec that parallel mtDNA and Y-chromosome inheritance. This allows detailed genealogical reconstruction up and down generations along maternal and paternal lines (17, 35, 36). We followed the fate of the genealogical lineages from the origins of the Quebec colony (1608) down to the years 1931-60. Using BALSAC, we provided a broader perspective on the creation of the Québec population than only using genetic data. This includes Québec population immigration history, well after the British Conquest in 1760, and factors and mechanisms underlying its demographic and geographical expansion.

Our study differs from previous analyses using ascending genealogies of existing populations (5, 35, 37), or the studies focused on descending data of a single lineage or limited to a region (16, 17, 38). It also differs from studies analyzing data sets from a limited number of parishes (12, 39), religion groups (40) or geographic isolates (10, 41). We used the BALSAC database that is unique in its completeness, quality, and breadth. It includes a quasi-complete reconstruction of the descending genealogies of Québec’s historical population of millions of individuals over more than a dozen generations. We know the founders whose marriages were likely to be registered and their descending marrieds recorded in parish documents of Québec. With this information in hand, we evaluated the yearly number of immigrants *N*_*i*_ and their contributing part *Nc*_*i*_, who left *Nw*_*i*_ carriers of their uniparental lines among 1931-60 married descendants. We introduced the survival ratio, SR_*i*_=*Nc*_*i*_/*N*_*i*_, to compare the success of consecutive waves of immigration that leave descendants (through maternal and paternal lines) among the 1931-60 married individuals. It is important not to confound SR_*i*_ with the wave *i* numerical outcome *Nw*_*i*_, i.e., the number of descendant copies among 1931-60 marrieds (Figure 1C). We also followed the EFS along all maternal and paternal lines. To tell apart the role of fertility, EFS, from extinction (due to drift or to emigration), we introduced additional descriptive measures. These are the net growth *dc*_*i*_ and the net extinction rate, *a*_*i*_. They estimate net growth or net extinction of distinct lines, respectively, between a time point *i* in the past and an extant population, such as that of the 1931-60 Québec (Figure 1E and F). We quantified the same demographic parameters for all QB as well.

We observed that during most of the history of Québec, the SR_*i*_ of the immigrants and QB individuals was declining (Figure 1D). SR_*i*_ represents the proportion of the contributing lineages. At first glance, while reasoning in terms of a simple demographic model of a population of a constant size, we found this time dependence of SR_*i*_ striking. Under genetic drift and in a growing population, SR_*i*_ is thought to increase with time. Genetic drift (42) is more potent when a population is small (43) and is thus expected to reduce the number of lineages more effectively at the start of a colony. In general, genetic drift gets weaker in a growing population (44, 45), which implies that the proportion of surviving lineages introduced closer to the present should be greater than that of the lineages introduced earlier. However, we observed the opposite, although perhaps not surprisingly so. As Alan Fix (3), while paraphrasing James Neel (2), stated: “the history of the human species and particular populations suggests that long-term equilibria, an assumption of many genetic models, is only a convenient mathematical fiction.” Different demographic and historical factors need to be considered to understand the genetic structure of extant populations (46).

### Effective family size and Québec evolutionary dynamics

Our study finds that the effective family size, EFS, remains a major player in the evolution of human populations. EFS represents the average number of children likely to increase offspring propagating generations down. The SR_*i*_ declines in lockstep with a decreased EFS (Figures 2 and 3), revealing a very close correlation between the EFS and their SR_*i*_ (Figure 3A and B). This correlation remains straightforward from the colony start until the end of the 19^th^-century: EFS determines the demographic outcome for many generations ahead.

The correlation between EFS and SR_*i*_ is not fortuitous. It reflects the fate of the introduced lineages that can be readily followed in the *dc*_*i*_ and *a*_*i*_ plots (Figures 1E and 1F). These plots reveal the interplay between prosperous and extinct lineages, respectively. The immigrants *a*_*i*_ soars dramatically at the turn of the 19^th^ century, whereas their *dc*_*i*_ drops (Figure 1E). The immigrants become less and less successful during the early 19^th^ century, and their number *N*_*i*_ eventually exceeds that of their 1931-60 descendants (*Nw*_*i*_). This starts when the *a*_*i*_ and *dc*_*i*_ plots cross when *Nw*_*i*_ = *Nc*_*i*_. After 1850, immigrants’ *dc*_*i*_ increases and *a*_*i*_ plateaus, but the balance is negative given *N*_*i*_*>Nw*_*i*_ (Figure 1E). Among QB, there is less variance in the corresponding plots (Figure 1F). QB *dc*_*i*_ falls at the turn of the 18^th^ century (matching that of the immigrants), levels off after this time and eventually rises during the 19^th^ century (Figure 1F). QB *a*_*i*_ never tops QB *dc*_*i*_, yet the portion of their extinct lineages steadily rises, with *a*_*i*_ noticeably increasing by the end of the 18^th^-century (Figure 1F, but see also Figure 3C and D).

Favored were immigrant parents who left plentiful married progeny. It puts EFS, a proxy of fitness, as the most important variable determining parental demographic success through future generations. However, for this success to persists, it requires the QB progeny to continue to be reproductively successful as overall SR is ultimately a function of successive EFS along descendant lines. And indeed, a correlation of family size between generations was described in various populations (5, 11, 27). Evidence also accumulates that reproductive success might be heritable, as shown in Québec and other preindustrial societies (Milot et al. 2011, Moreau et al. 2011, Pettay et al. 2005, Stearns et al. 2010). This reproductive success could have a genetic basis. It might imply local selection (46) by acting on new mutations and/or standing variation (47-49). High EFS creates a surplus of new combinations among existing variants and increases new haplotypic arrangements. Likewise, non-recombining haplotypes of mtDNA and Y-chromosome could partake in these new combinations. Some of these genetic changes may present increased adaptive values. High EFS, continued over many generations given their high SR, suggests that some of these combinations were demographically favored. By demographic favor/gain, we understand biological (evolutionary), social, cultural, and economic benefits conducive to population expansion. In Québec, despite harsh environmental conditions, it seems to include many winning factors supporting survival.

### Factors affecting EFS estimates in Québec

We cannot ignore couples who failed to have children for various biological reasons, such as in obvious cases when one of the partners was sterile (50, 51). Sterility affects about 15% of contemporary couples (52) and was estimated to be about 10% in historical times. Therefore, the sterility of one of the partners must have contributed to the “zero fertility” peak in plots of the number of effective children that was published previously (16, 27). The overall childlessness reported in our plots corresponds to a cumulative effect of biological (see above) and other reasons that results in effective children being absent in the BALSAC records. If no married progeny is found in the BALSAC database, it can mean that the parents did have children, but these children died before reaching reproductive age; they did not marry or moved outside of Québec (25, 53). If the childlessness is only due to biological causes, we would expect this to be relatively even among different Québec regions. However, we observed that zero peaks (no children among married couples) markedly differed among the regional populations and often, this difference exceeded10-15% (Figure S9).

Furthermore, the proportion of parents with no recorded children is more significant among immigrants than among QB (Figure S10). This is consistent with the emigration of the first-generation immigrants or their children, who moved out of Québec (54) and disappeared from the BALSAC horizon (Figure 1E and F). This already occurred in the early colonial times when some immigrant settlers moved outside Québec borders and moved southwards to the United States and western Canada (25, 55). This phenomenon became exceptionally substantial in the 19^th^-century, engulfing many QB (53-55). Emigration was motivated by unsatisfactory local conditions and tempting opportunities to prosper elsewhere. Local economic and social conditions started to worsen due to the rise in population density, military conflicts, famines, and epidemics (25, 30, 56, 57). This emigration affected the newcomers considerably more than the previous settlers. Therefore, when immigrants and their children left Québec, then “lost or extinct” does not necessarily imply that lineages absent from BALSAC records did not prosper elsewhere. In our calculations, we included only those that remained within the Québec borders. However, when necessary, we used BALSAC data from outside Québec to identify common maternal or paternal founders among the immigrants.

Québec population was not the first colony benefitting from over-Atlantic European expansion. Europeans re-colonized different areas of the globe in historical times, often founding new nation-states, currently populated with their descendants, with various demographic and genetic outcomes. In Meso- and South America, a skewed sex ratio among European invaders, led to an asymmetric male-biased admixture with Native populations. Up to today, European Y-chromosomes and Native mtDNA prevail in the descendant populations (58-61). Also, in Quebec, during the 17^th^-century colonization of the St. Lawrence Valley by the French, male settlers were more numerous than female immigrants (30). Genetic studies indicate that at that time, admixture with Natives was limited (8). Also, according to Catholic Church records, only a few Native females contributed to the French-Canadian genealogy (31, 35, 62). BALSAC database covers most of Québec genealogies, but not all of the Québec diversity. Missing information includes some unnoted Natives and many Protestant Church records.

### Québec population history and founder events

In genetic studies of the Québec population, it was usually presumed that French-Canadians descended from about 8,500 immigrants, (15) who settled in Nouvelle France before the British conquest in 1760 (e.g., (7, 27, 29, 30, 37). We found that this premise is only partially correct. We show (Figures 1 and 2; Tables 1 and 2) that immigrants after the 1750s added a multitude of new matrilineal and patrilinear lineages that were still present in the 1931-60 genetic pool. They exceeded the contribution of the first settlers in terms of the number of new lineages but contributed much less to the mass of the genetic makeup of the post-1930 population (Table 2; Figure S5). And indeed, there is a striking difference between the 17^th^ and the early 19^th^-century immigration. The population of the first immigrants was thrifty and thriving, populating a sparsely inhabited territory. In contrast, the subsequent immigrants were numerous, but their immigration success was declining: the territory was already packed, and many immigrants failed to settle. It could be that during the 17^th^ century, there was more social and economic cooperation, despite men’s rivalry to find a spouse (15, 30). Although there were local conflicts with the Natives and the British, the environment was challenging but healthy since epidemics arrived only by the late 17^th^-century (57). Therefore, being first and successful would have given these individuals the edge over subsequent generations in a quickly growing population (16, 63, 64). Québec’s example confirms this view. In the early 19^th^ century, Québec was already occupied by the well-established and prospering QB individuals. For newcomers, a suitable economic niche was more challenging to find. It persuaded many immigrants to settle in Québec only temporarily and eventually move elsewhere.

Consequently, their EFS was dropping. Also, at that time, QB families were affected, albeit less dramatically. The fraction of reproductively prosperous families decreased (Figure 2). Interestingly, however, this occurred not everywhere, since the late 19^th^-century population of Saguenay–Lac-Saint-Jean was demographically as successful as that of the 17^th^ century Québec settlers (16, 65).

Québec colonization can be considered as a series of successive founder events. It was accompanied by the founding of genetic mutations causing regionally distributed rare and endemic hereditary diseases (“medical founder effects), well-documented in the Saguenay-Lac-St.-Jean and Charlevoix regions (65-68). However, besides this medical interest, Québec also provides a founder population model in terms of the original concept developed by Ernst Mayr (69). He stated that a relatively small group of migrants establishes a new population. Due to a sampling process, the overall diversity was reduced, and many variants were moved to different classes of the allelic frequency spectrum. Likewise, while many rare variants were left behind, some, including defective, disease-causing alleles, could locally increase in frequency, thereby explaining “medical founder effects” (66, 70, 71). As shown here, in Quebec, there were not punctual founder events, such as occurred when a foreign immigrant species accidentally seed a remote island and then evolves in isolation (72). As often was the case in human migrations, founder populations remained in contact with forefront groups that colonized new territories. Here specifically, familial bounds led to non-random migration of lineages so that an individual would leave France to immigrate to Québec where his sibling had already settled (73). Another example is when deported Acadians returned to New France (55). Such immigration pattern is typical to expanding human populations settling in an empty territory or conquering already occupied areas (74, 75).

At an evolutionary scale, human populations went through expansions followed by contractions and sometimes near extinctions (4) to eventually recover after moving to favorable environments conducive to growth, development, and range expansion. The “out-of-Africa” migration was modeled as a serial founder effect, where migrating groups went through a series of bottlenecks accompanied by range expansions followed by demographic growth. This migration gradually reduced the genetic diversity of these populations because of their distance from Africa (76). At the micro-evolutionary scale of fewer than 20 generations, Quebec demography may serve as an example of how human populations evolved, assisting interpretation of models of demographic inference attempting to reconstruct demographic histories (1).

Continental dispersals occurred multiple times in the history of the human species (77). The migratory history of *Homo sapiens* can be traced through the worldwide geographic distribution of its genetic variants (75, 78-80). It also reveals past evolutionary processes, including a positive and purifying selection that adapted settlers and their descendants to local environments (81, 82). In that respect, recent demographic colonisations and expansions by humans (i.e., in the last hundreds to thousands of years) are thought to have contributed to the build-up of the genetic burden and architecture of diseases in contemporary populations through founder effects and relaxation of purifying selection against harmful alleles (68, 83-85). However, this hypothesis remains challenged by mixed empirical support and theoretical considerations (44, 86-88). For example, earlier immigrants benefited from demographic gain/advantage reproducing when the growing population was small. They so boosted their genetic representation in future generations relative to latecomers (38, 63, 89, 90). The earliest lineages can also benefit from their seniority because they are ahead of subsequent immigration waves in adapting to their new environment through the plasticity of evolutionary changes. As shown here, founder effects could have played a crucial role in the evolution of the human species by providing conducive conditions for selection under the new environment and high EFS.

## Acknowledgments

We are indebted to Claudia Moreau and Jean-Francois Lefebvre, who conducted initial analyses of the BALSAC data, and especially to Eef Harmsen, as well as to Jerzy Kolasa and Brad Loewen, for their comments on the manuscript. We are also thankful to members of the BALSAC team for their help in exploring the database. Computations were made on the Cedar and Mammouth supercomputers at Simon Fraser University and the Université de Sherbrooke, respectively, managed by Compute Canada and Calcul Québec, funded by the Canada Foundation for Innovation, the Ministère de l’Économie et de l’Innovation du Québec and the Fonds de recherche du Québec – Nature et technologies. This research was supported by the Fonds de recherche du Québec – Santé, via the Réseau de médecine génétique appliquée (DL, HV), the National Sciences and Engineering Research Council of Canada (DL), and by DL donations via la Fondation de l’Hôpital Sainte-Justine.

---

**Figure S1.**
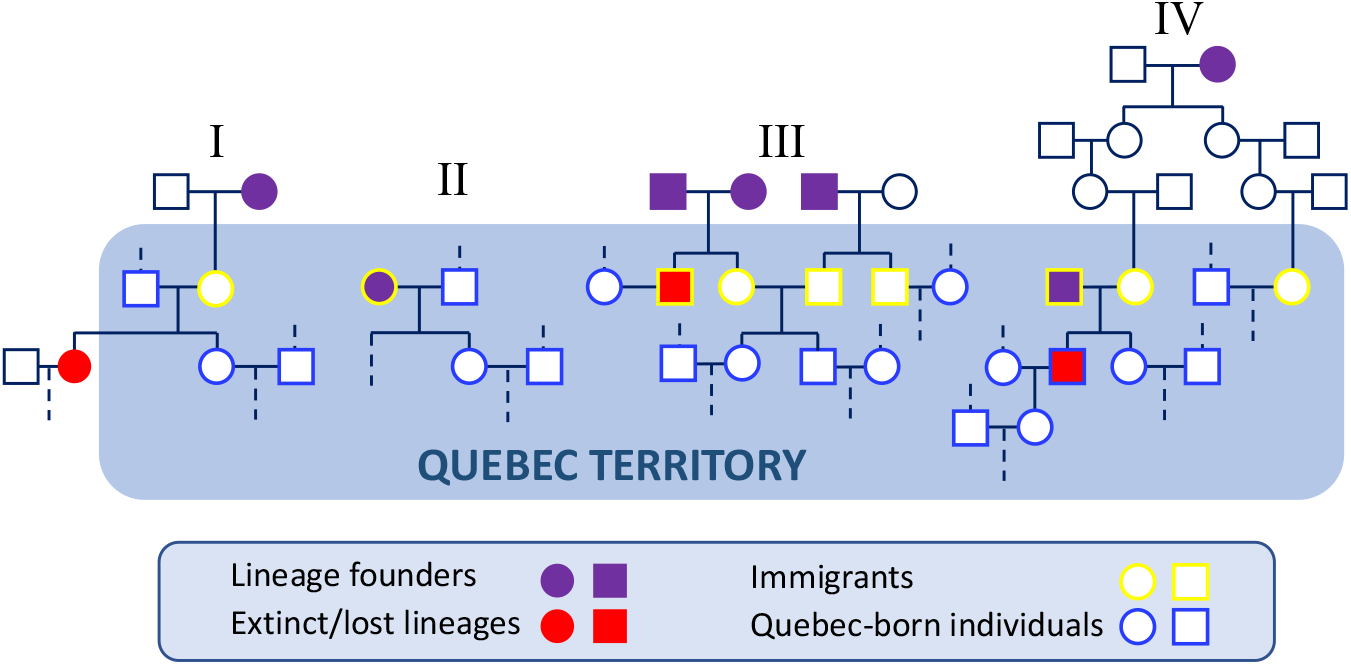
An example of matrilineal and patrilineal founders, immigrants, and Quebec-born individuals. We define immigrants as the first generation of settlers to marry in Quebec (symbols circled in yellow; maternal immigrant in example I). Although strictly speaking, some of these immigrants were born in Quebec to immigrant parents. Immigrants can be lineage founders (symbols circled in a yellow frame filled with violet, example II). Genealogically recorded non-immigrant lineage founders are marked in violet. If their immigrant progenitors are the sole introducers of these lineages, as shown in example I. Their genealogical impact is equivalent to that of solitary immigrants, as shown in example II. Lineage founders who are ancestors of more than one immigrant represent the same lineage introduced more than once. This may be the case of siblings who immigrated together (example III) or with distantly related cousins (example IV). Whenever the genealogical information permits, we avoid counting more than once the same patrilineal or matrilineal lineage introduced by immigrant siblings or immigrant cousins. Some immigrant lineages do not contribute to present-day Quebec diversity due to extinction by emigration (example I); by lack of daughters and/or sons altogether (examples III and IV, red-filled symbols).

**Figure S2.**
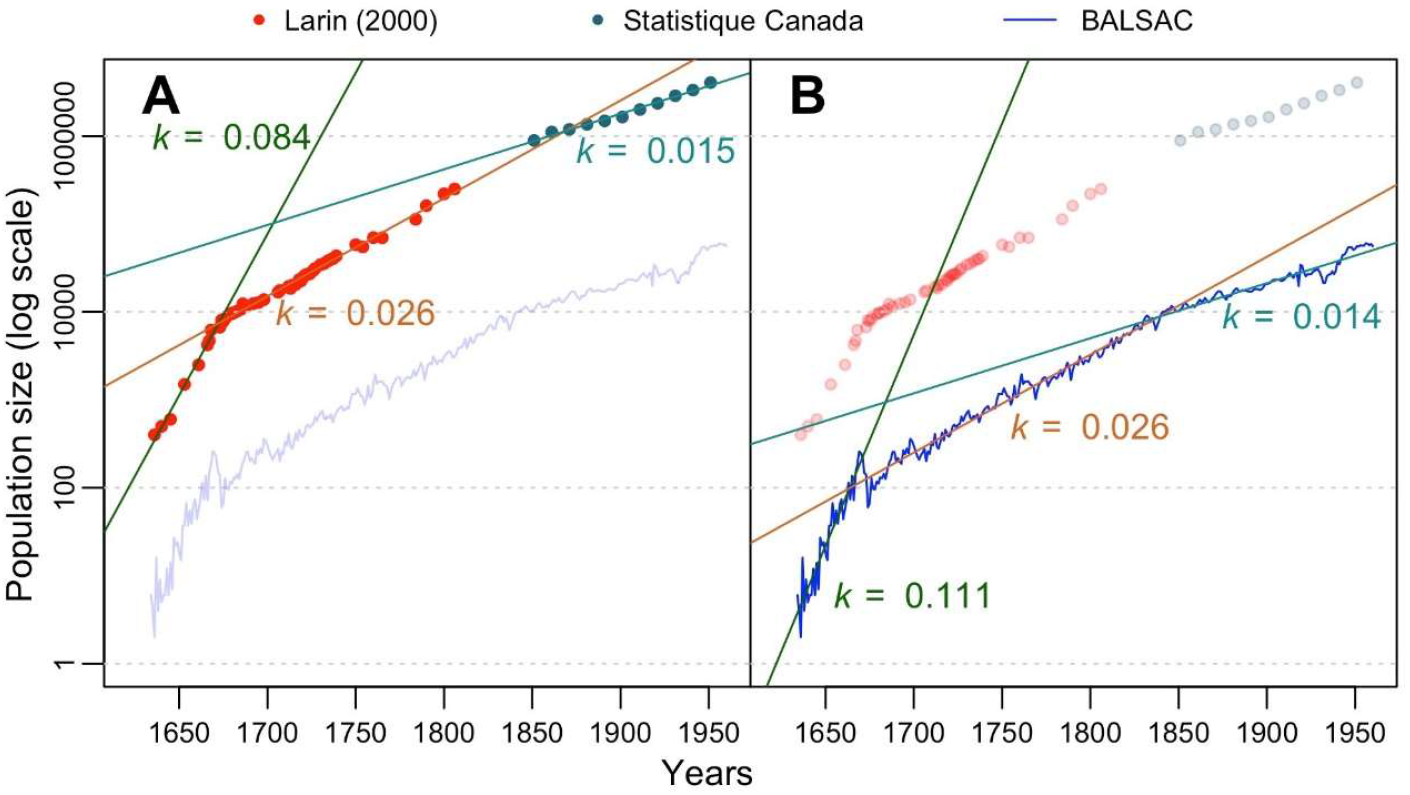
Estimates of the Quebec population growth rate. Growth rate (*k*) is estimated from the slope of the plot of historical population size *versus* time. In (**A**), we used population size data reported by Larin (red circles) ((25) and references therein) and Statistics Canada census data since 1851 (green circles) (http://www.stat.gouv.qc.ca/default_an.html). In (**B**), to estimate *k*, we used the yearly number of BALSAC recorded marriages (blue line – shown in pale blue in (**A**)) as a proxy for the population size. Linear regression curves were calculated by fitting the log of population sizes to the years by periods: 1621-1670 in dark green, 1671-1850 in brown and 1851-1960 in cyan. The associated slope coefficients (*k*) are displayed using the same color code.

**Figure S3.**
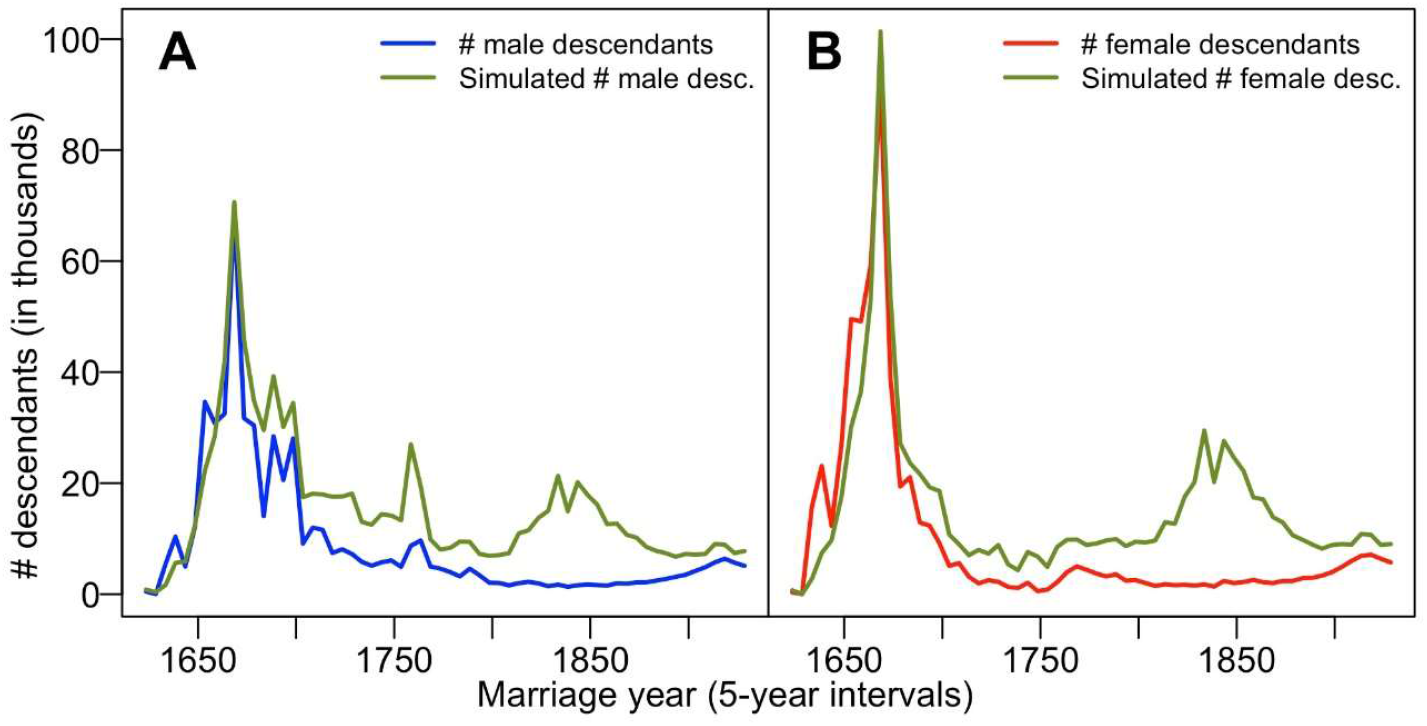
Actual numbers of 1931-60 descendants from immigration waves since 1621 compared to their simulated numbers assuming a constant survival ratio of 0.45 and 0.55, respectively, and constant annual growth rate (*k*) of 0.021 and 0.022 male and female lineages, respectively. In (**A**), paternal lineages with the observed (blue) and simulated (green) numbers of male descendants. (**B**) Maternal lines with the observed (red) and simulated (green) numbers of female descendants.

**Figure S4.**
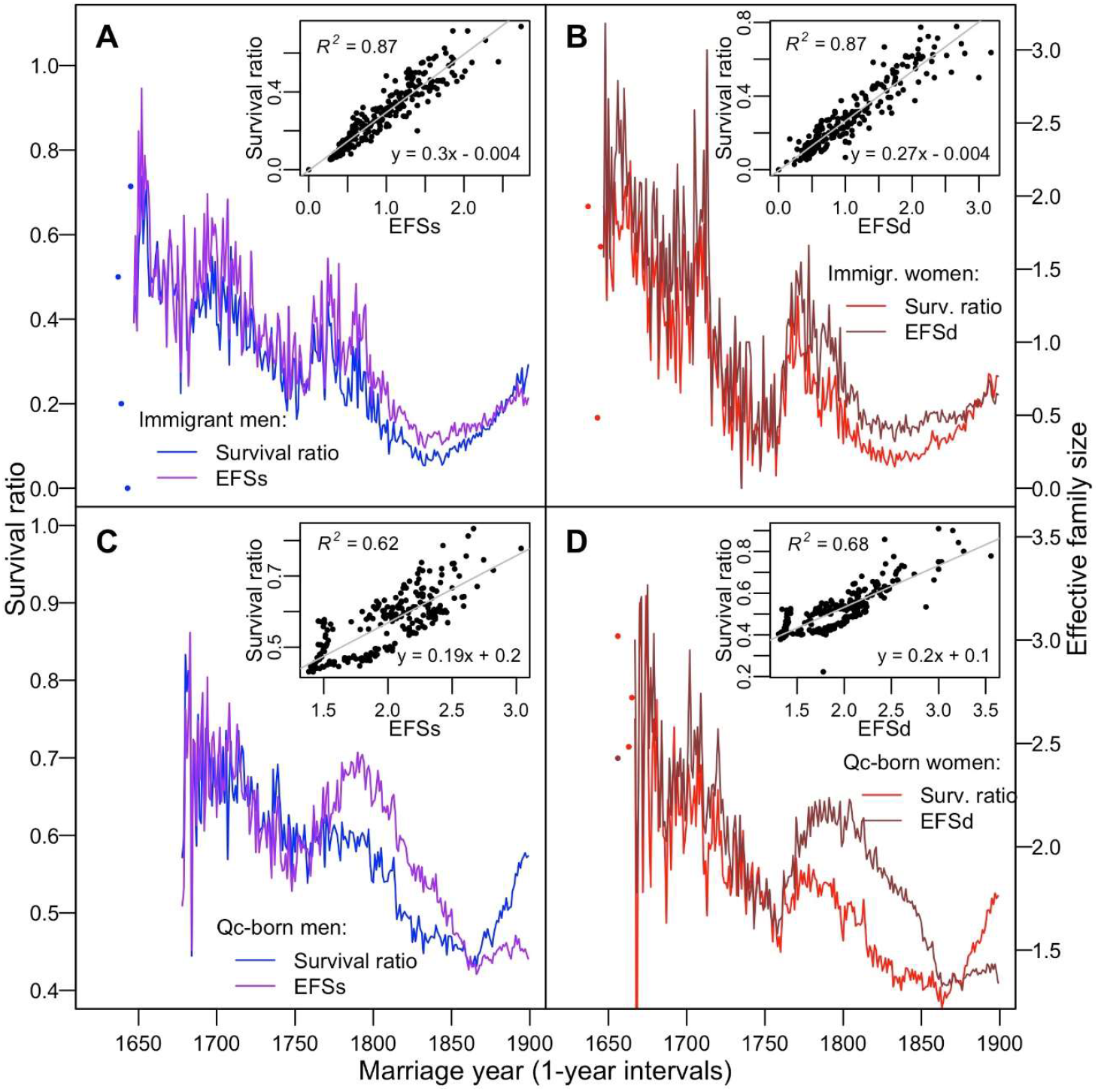
Correlation between the effective family size and the survival ratio, as in Figure 3 in the main text, here based on 1-year intervals. Overlapping plots of total EFSs and EFSd and their survival ratios for the immigrants (**A** and **B**) and the Quebec-born lineages (**C** and **D**). Their correlation plots are shown in the insets.

**Figure S5.**
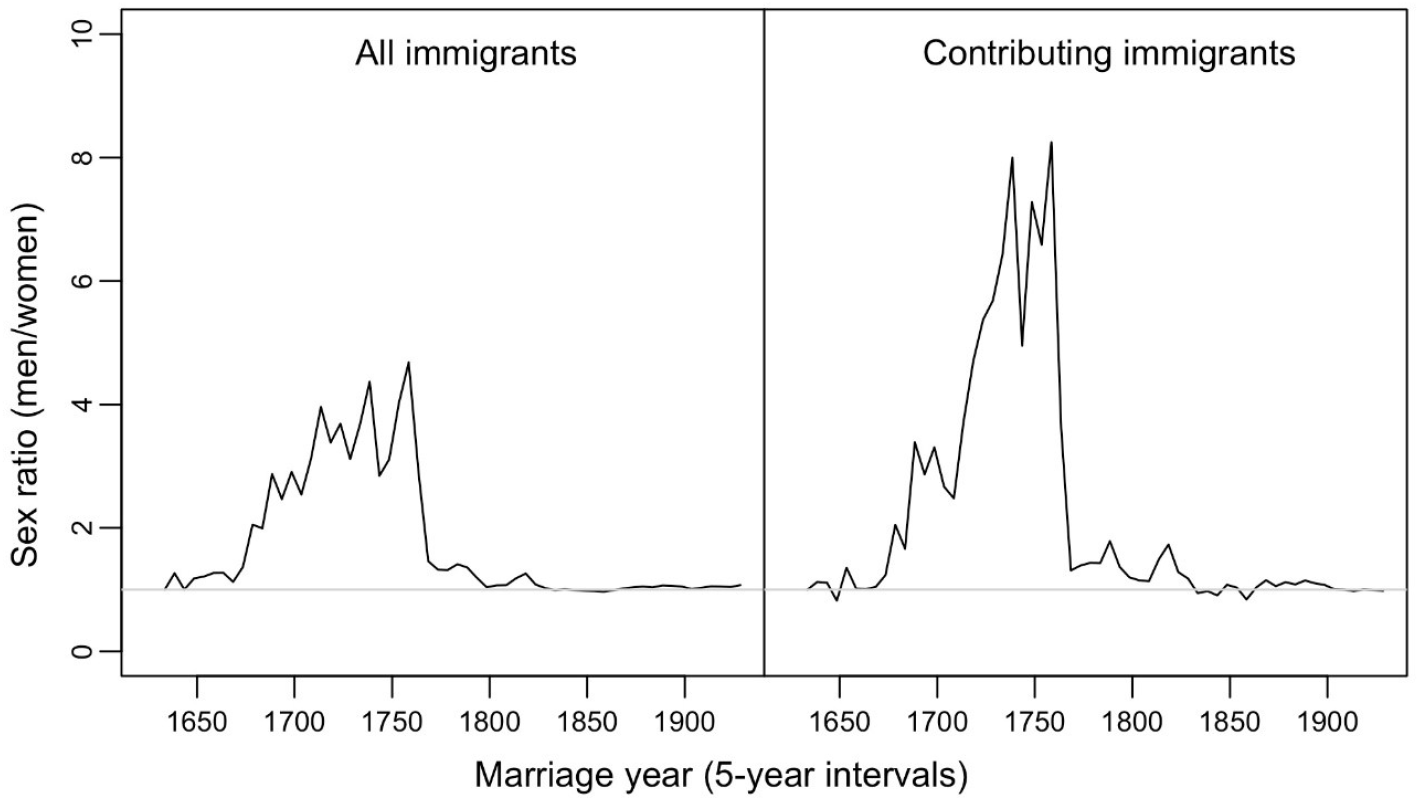
Man to women sex ratio among the immigrants. The number of male immigrants divided by the number of female immigrants considering all immigrants (**left panel**) and only a subset of the contributing ones (**right panel**).

**Figure S6.**
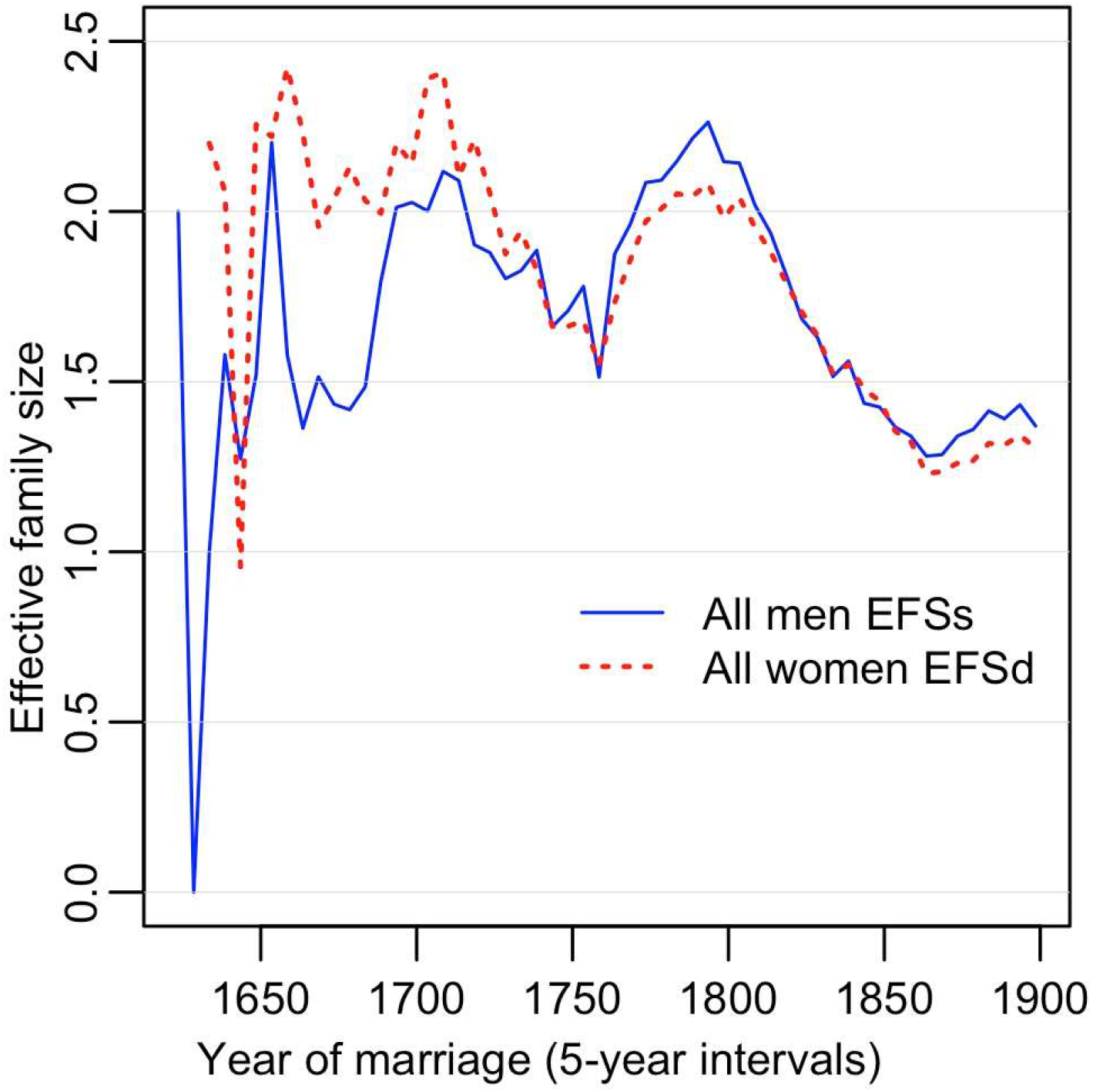
Comparing EFS-sons (EFSs) and EFS-daughters (EFSd) of all marrieds. Please, note much lower EFS-sons (EFSs) than that of EFS-daughters (EFSd) before 1700, at the second half of the 17^th^ century. This is consistent with the scenario of many Québec-born men, from immigrant and non-immigrant parents at that time, to leave Québec exploring other territories referred to as Nouvelle France.

**Figure S7.**
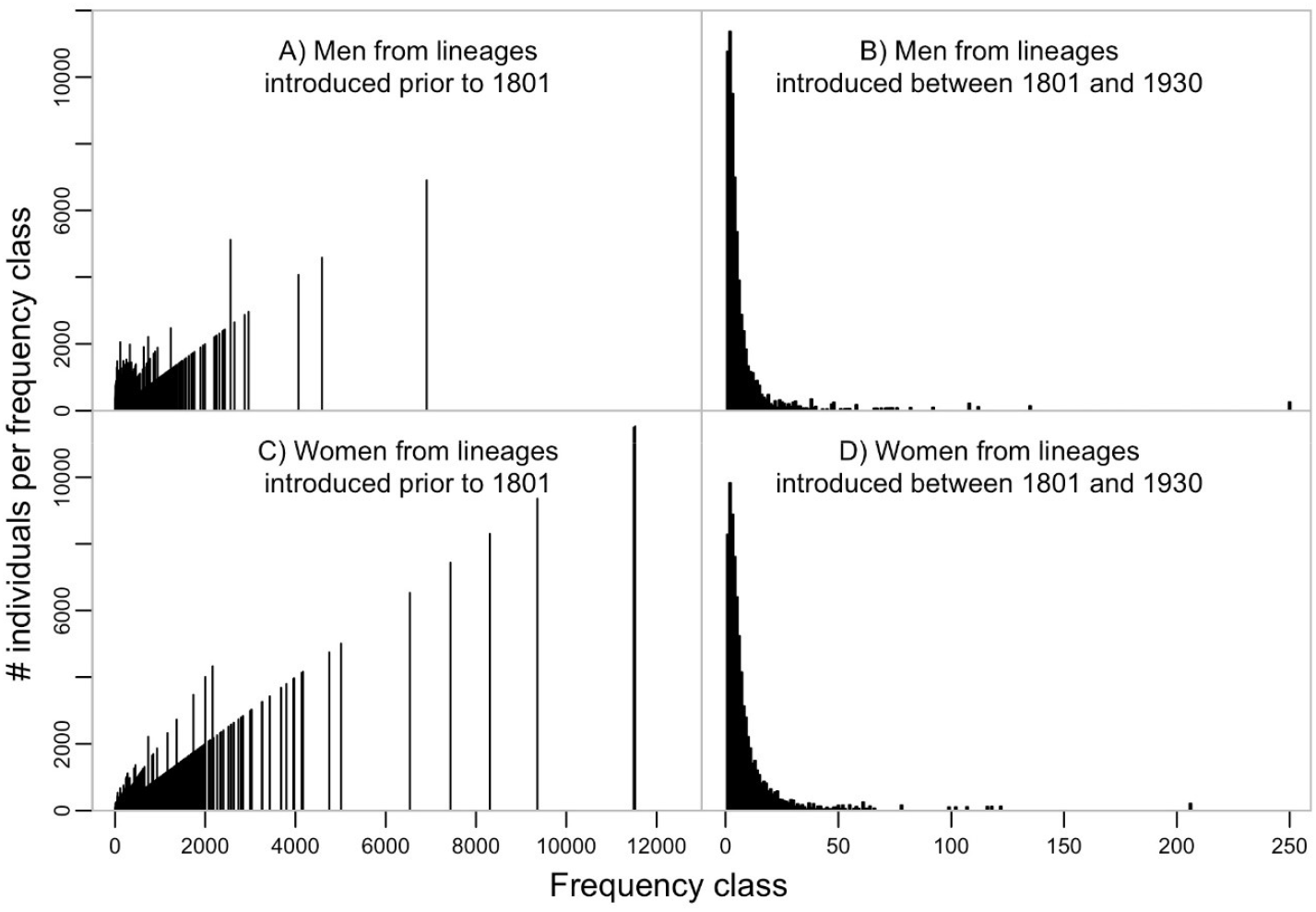
Frequency spectra of maternal and paternal lineages from before 1931 in the 1931-60 population. The lineages introduced before 1801 are shown in the left panels, and those introduced between 1801 and 1930 are in the right plots. Note that on the y-axis, we present the whole number of individuals within the frequency class to make the plot more informative and transparent as in (31). A typical plot of frequency classes registers only the number of classes on the y-axis. As a result, frequency classes represented only a few times or once, practically disappear from graphic representation, especially when frequency classes on the left are particularly numerous.

**Figure S8.**
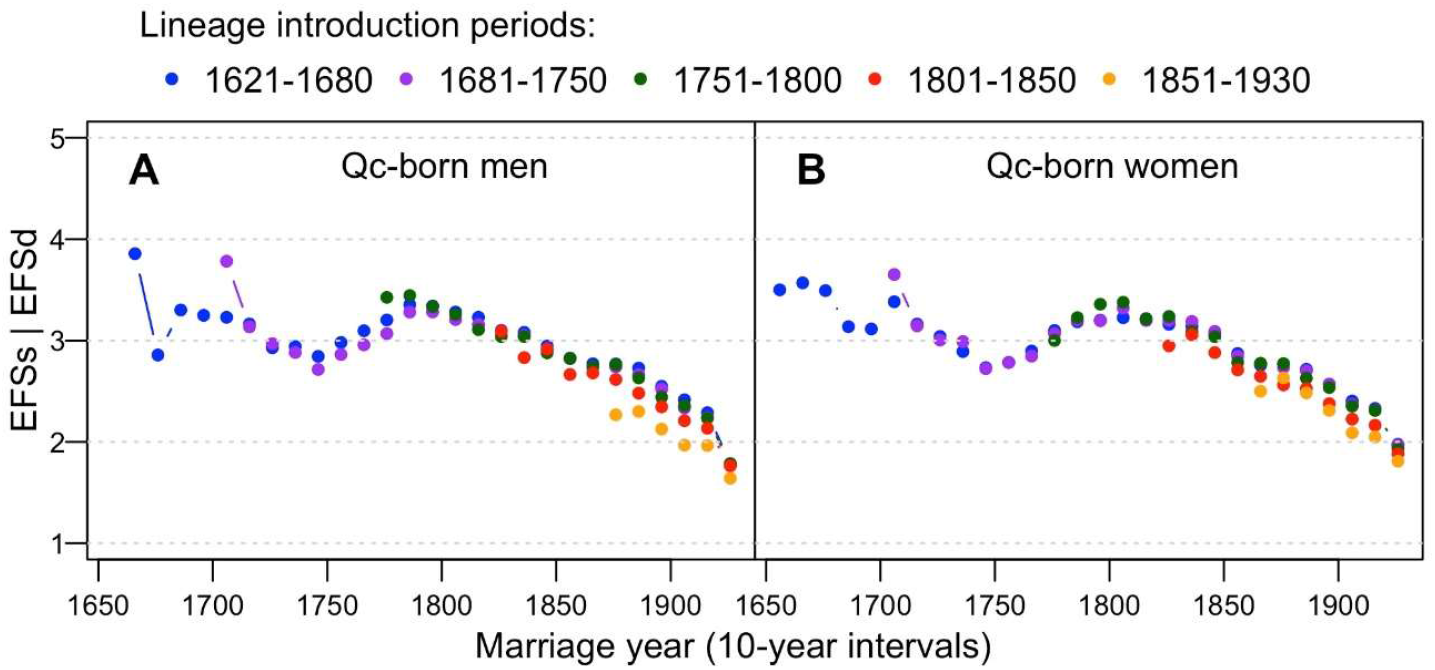
EFS-sons and EFS-daughters of the contributing lineages introduced at different periods. EFS is averaged over 10-year intervals, and progeny of lineages at different periods are marked by different colors as indicated. Remember that contributing refers to lineages that are still present within the 1931-60 Québec population.

**Figure S9.**
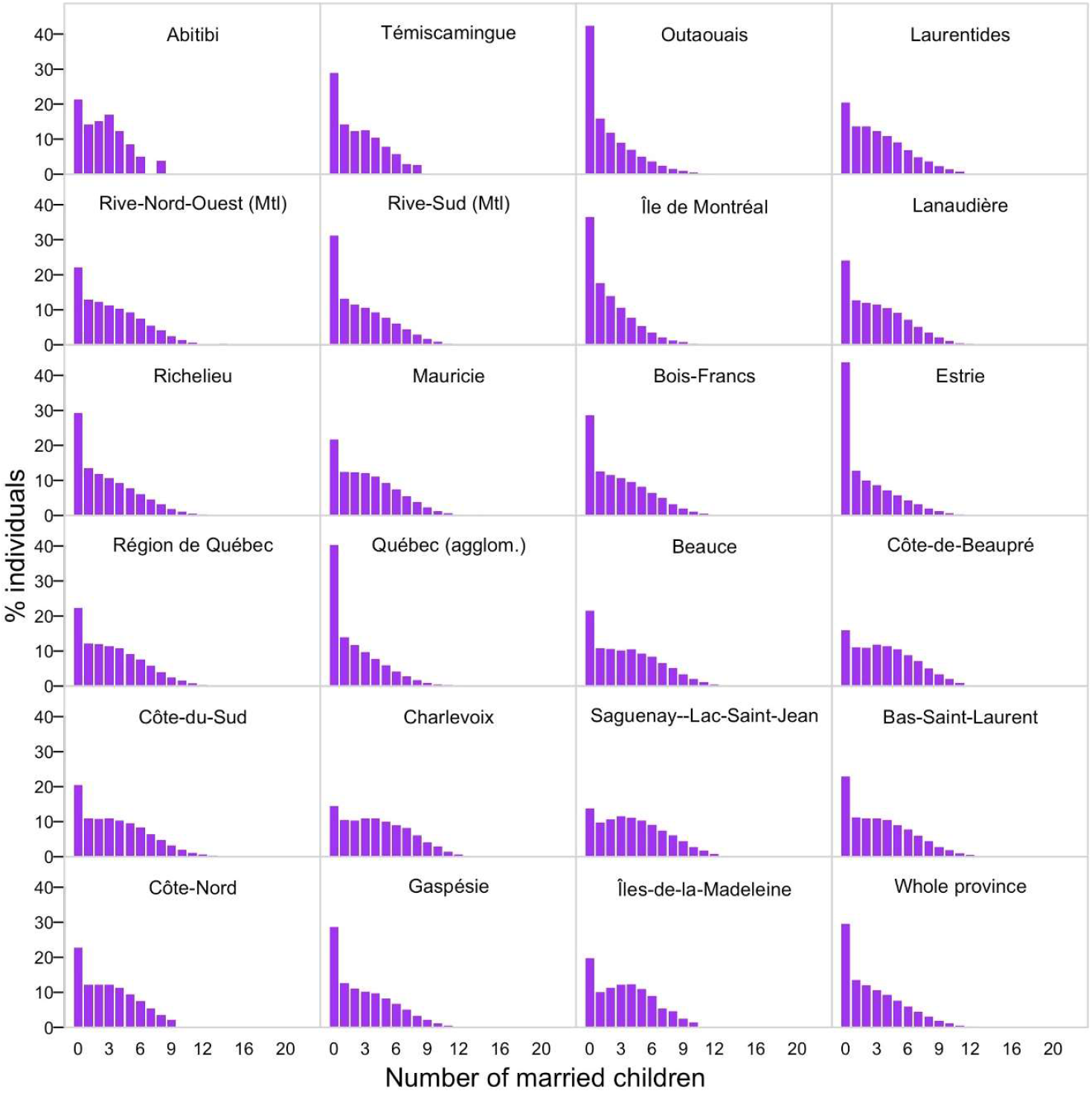
Distribution of the number of married children (total EFS) in families from different regions of Quebec. Because only children married in Québec (registered in BALSAC) are counted, it emphasizes the potential effect of emigration out of Québec on the regional EFS frequency distribution. Note that “zero frequency EFS” exceeds the 10-15% threshold well, ascribed to couple sterility in many regions. Likewise, convex histograms of subsequent frequency classes, such as in Charlevoix, for example, become concave when preceded by high zero EFS frequency, as in Outaouais or Isle de Montréal. Emigration affects the EFS of all families. Besides emigration (strictly, leaving Québec and married or not elsewhere), other factors: economic, social, and historical (e.g., wars, epidemics) could have reduced BALSAC recorded EFS.

**Figure S10.**
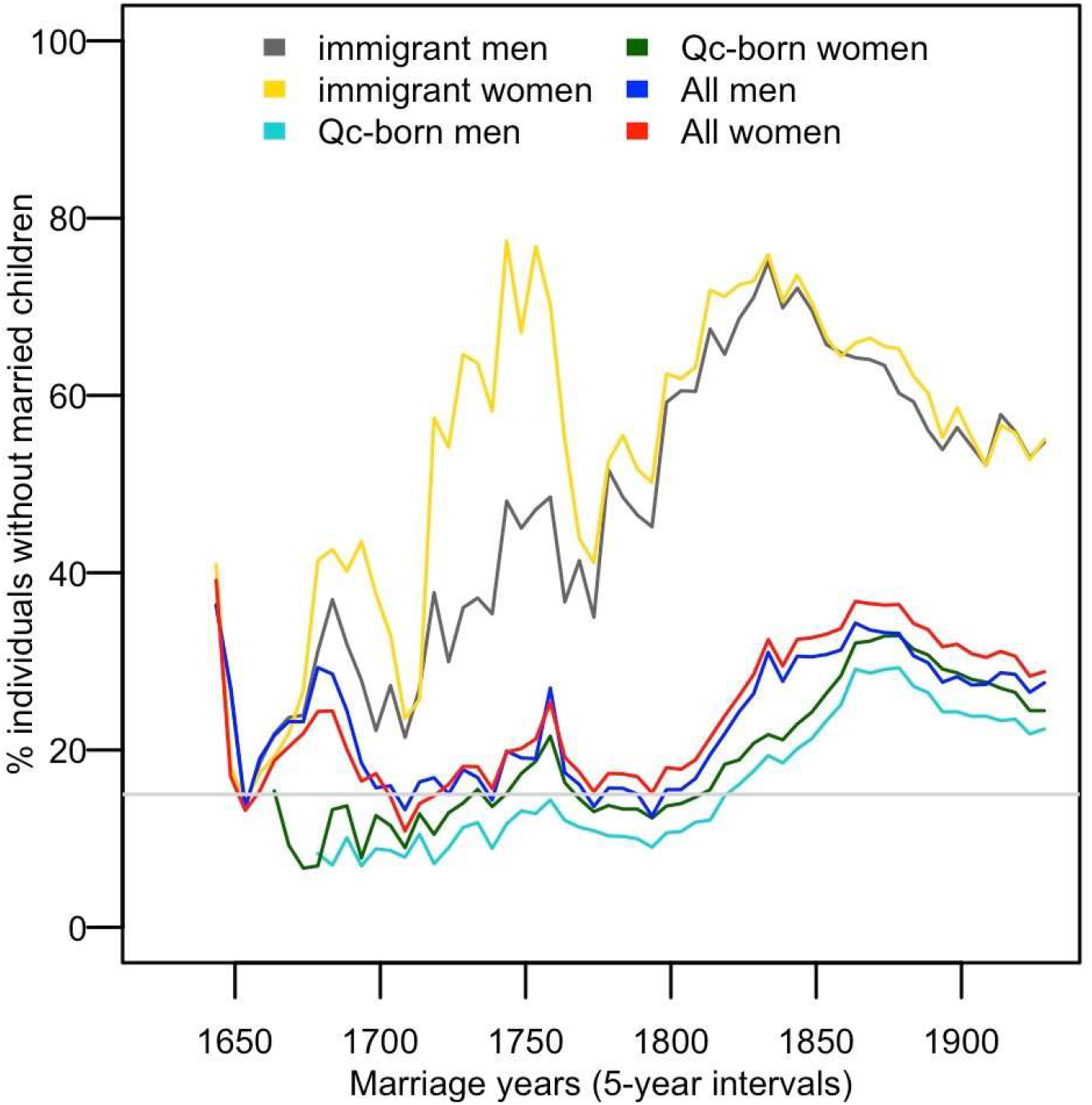
Proportion of different categories of parents with no married children, i.e., of zero EFS. Percentages of immigrant men (dark grey line), immigrant women (yellow line), QB men (cyan line), QB women (green line), all men (blue line), and all women (red line) without married children as a function of their marriage year by 5-year intervals. The light grey line indicates an estimate of the infertility rate (15%; (52)).

## References

1. Marchi N, Schlichta F, Excoffier L. Demographic inference. Curr Biol. 2021;31(6):R276–R9.

2. Neel JV. The circumstances of human evolution. Johns Hopkins Med J. 1976;138(6):233–44.

3. Fix AG. ANTHROPOLOGICAL GENETICS OF SMALL POPULATIONS. Annual Review of Anthropology. 1979;8:207–30.

4. Takahata N. Repeated failures that led to the eventual success in human evolution. Mol Biol Evol. 1994;11(5):803–5.

5. Helgason A, Hrafnkelsson B, Gulcher JR, Ward R, Stefansson K. A populationwide coalescent analysis of Icelandic matrilineal and patrilineal genealogies: evidence for a faster evolutionary rate of mtDNA lineages than Y chromosomes. Am J Hum Genet. 2003;72(6):1370–88.

6. Ballantyne KN, Goedbloed M, Fang R, Schaap O, Lao O, Wollstein A, et al. Mutability of Y-chromosomal microsatellites: rates, characteristics, molecular bases, and forensic implications. Am J Hum Genet. 2010;87(3):341–53.

7. Roy-Gagnon MH, Moreau C, Bherer C, St-Onge P, Sinnett D, Laprise C, et al. Genomic and genealogical investigation of the French Canadian founder population structure. Hum Genet. 2011;129(5):521–31.

8. Moreau C, Lefebvre JF, Jomphe M, Bherer C, Ruiz-Linares A, Vezina H, et al. Native American admixture in the Quebec founder population. PLoS One. 2013;8(6):e65507.

9. Jonsson H, Sulem P, Kehr B, Kristmundsdottir S, Zink F, Hjartarson E, et al. Parental influence on human germline de novo mutations in 1,548 trios from Iceland. Nature. 2017;549(7673):519–22.

10. Roberts DF. Genetic Effects of Population Size Reduction. Nature. 1968;220:1084.

11. Pluzhnikov A, Nolan DK, Tan Z, McPeek MS, Ober C. Correlation of intergenerational family sizes suggests a genetic component of reproductive fitness. Am J Hum Genet. 2007;81(1):165–9.

12. Pettay JE, Kruuk LE, Jokela J, Lummaa V. Heritability and genetic constraints of life-history trait evolution in preindustrial humans. Proc Natl Acad Sci U S A. 2005;102(8):2838–43.

13. Morera B, Barrantes R. Is the Central Valley of Costa Rica a genetic isolate? Rev Biol Trop. 2004;52(3):629–44.

14. BALSAC. BALSAC population register website. [Available from: http://balsac.uqac.ca/.

15. Charbonneau H, Desjardins, B., Guillemette, A., Landry, Y., Légaré, J., Nault, F. Naissance d’une population. Les Français établis au Canada au XVIIe siècle: Institut National d’Études Démographiques, Presses de l’Univéersité de Montréal, Presses Universitaires de France; 1987. 232 p.

16. Moreau C, Bherer C, Vezina H, Jomphe M, Labuda D, Excoffier L. Deep human genealogies reveal a selective advantage to be on an expanding wave front. Science. 2011;334(6059):1148–50.

17. Milot E, Moreau C, Gagnon A, Cohen AA, Brais B, Labuda D. Mother’s curse neutralizes natural selection against a human genetic disease over three centuries. Nat Ecol Evol. 2017;1(9):1400–6.

18. Vezina H, Durocher F, Dumont M, Houde L, Szabo C, Tranchant M, et al. Molecular and genealogical characterization of the R1443X BRCA1 mutation in high-risk French-Canadian breast/ovarian cancer families. Hum Genet. 2005;117(2-3):119–32.

19. Nelson D, Moreau C, de Vriendt M, Zeng Y, Preuss C, Vezina H, et al. Inferring Transmission Histories of Rare Alleles in Population-Scale Genealogies. Am J Hum Genet. 2018;103(6):893–906.

20. Salem AH, Badr FM, Gaballah MF, Paabo S. The genetics of traditional living: Y-chromosomal and mitochondrial lineages in the Sinai Peninsula. Am J Hum Genet. 1996;59(3):741–3.

21. Lippold S, Xu H, Ko A, Li M, Renaud G, Butthof A, et al. Human paternal and maternal demographic histories: insights from high-resolution Y chromosome and mtDNA sequences. Investig Genet. 2014;5:13.

22. Légaré J. A population register for Canada under the French Regime: context, scope, content and appplications. Canadian Studies in Population. 1988;15(1):1–16.

23. Vézina H, Bournival JS. An overview of the BALSAC database: past developments, current state and future prospects. Historical Life Course Studies. 2020;11(2):1–17.

24. Tremblay M, Vezina H. New estimates of intergenerational time intervals for the calculation of age and origins of mutations. Am J Hum Genet. 2000;66(2):651–8.

25. Larin R. Brève histoire du peuplement européen en Nouvelle-France. Sillery (Québec): Septentrion; 2000. 226 p.

26. StatisticsCanada. https://web.archive.org/web/20080501112831/http://www40.statcan.ca/l01/cst01/demo62f.htm. 2005.

27. Austerlitz F, Heyer E. Social transmission of reproductive behavior increases frequency of inherited disorders in a young-expanding population. Proc Natl Acad Sci U S A. 1998;95(25):15140–4.

28. Thomson CE, Bayer F, Crouch N, Farrell S, Heap E, Mittell E, et al. Selection on parental performance opposes selection for larger body mass in a wild population of blue tits. Evolution. 2017;71(3):716–32.

29. Livi Bacci M. The Population of Europe. Malden, MA: Blackwells; 2000.

30. Charbonneau H, Desjardins B, Légaré J, Denis H. The population of the St-Lawrence Valley, 1608-1760. In: Haines MR, Steckel RH, editors. A population history of North America. New York: Cambridge University Press; 2000. p. 99–142.

31. Moreau C, Vezina H, Yotova V, Hamon R, de Knijff P, Sinnett D, et al. Genetic heterogeneity in regional populations of Quebec--parental lineages in the Gaspe Peninsula. Am J Phys Anthropol. 2009;139(4):512–22.

32. Tremblay M, Jomphe M, Vézina H. Au nom des pionnières, de leurs filles et de toute leur descendance. Histoire Québec. 2011;17:29–30.

33. Hein J, Schierup M, Wiuf C. Gene Genealogies, Variation and Evolution. A Primer in Coalescent Theory: Oxford University Press, Oxford; 2005.

34. Wakeley J. Coalescent Theory: An Introduction. Greenwood Village, Colorado: Roberts & Company Publishers; 2009.

35. Moreau C, Vezina H, Jomphe M, Lavoie EM, Roy-Gagnon MH, Labuda D. When genetics and genealogies tell different stories-maternal lineages in Gaspesia. Ann Hum Genet. 2011;75(2):247–54.

36. Harding T, Milot E, Moreau C, Lefebvre JF, Bournival JS, Vezina H, et al. Historical human remains identification through maternal and paternal genetic signatures in a founder population with extensive genealogical record. Am J Phys Anthropol. 2020;171(4):645–58.

37. Bherer C, Labuda D, Roy-Gagnon MH, Houde L, Tremblay M, Vezina H. Admixed ancestry and stratification of Quebec regional populations. Am J Phys Anthropol. 2011;144(3):432–41.

38. Milot E, Mayer FM, Nussey DH, Boisvert M, Pelletier F, Reale D. Evidence for evolution in response to natural selection in a contemporary human population. Proc Natl Acad Sci U S A. 2011;108(41):17040–5.

39. Helle S, Lummaa V, Jokela J. Marrying women 15 years younger maximized men’s evolutionary fitness in historical Sami. Biol Lett. 2008;4(1):75–7.

40. Pichler I, Fuchsberger C, Platzer C, Caliskan M, Marroni F, Pramstaller PP, et al. Drawing the history of the Hutterite population on a genetic landscape: inference from Y-chromosome and mtDNA genotypes. Eur J Hum Genet. 2010;18(4):463–70.

41. Soodyall H, Nebel A, Morar B, Jenkins T. Genealogy and genes: tracing the founding fathers of Tristan da Cunha. Eur J Hum Genet. 2003;11(9):705–9.

42. Masel J. Genetic drift. Curr Biol. 2011;21(20):R837–8.

43. Buri P. Gene frequency in small populations of mutant Drosophila. Evolution; international journal of organic evolution. 1956;10:367–402.

44. Tajima F. Statistical method for testing the neutral mutation hypothesis by DNA polymorphism. Genetics. 1989;123(3):585–95.

45. Keinan A, Clark AG. Recent explosive human population growth has resulted in an excess of rare genetic variants. Science. 2012;336(6082):740–3.

46. Gravel S. When Is Selection Effective? Genetics. 2016;203(1):451–62.

47. Barrett RD, Schluter D. Adaptation from standing genetic variation. Trends Ecol Evol. 2008;23(1):38–44.

48. Peter BM, Huerta-Sanchez E, Nielsen R. Distinguishing between selective sweeps from standing variation and from a de novo mutation. PLoS Genet. 2012;8(10):e1003011.

49. McCoy RC, Akey JM. Selection plays the hand it was dealt: evidence that human adaptation commonly targets standing genetic variation. Genome Biol. 2017;18(1):139.

50. Mascarenhas MN, Flaxman SR, Boerma T, Vanderpoel S, Stevens GA. National, regional, and global trends in infertility prevalence since 1990: a systematic analysis of 277 health surveys. PLoS Med. 2012;9(12):e1001356.

51. Krausz C. Editorial for the special issue on the molecular genetics of male infertility. Hum Genet. 2021;140(1):1–5.

52. Matzuk MM, Lamb DJ. Genetic dissection of mammalian fertility pathways. Nat Cell Biol. 2002;4 Suppl:s41–9.

53. Vermette D. A distinct alien race. The untold story of Franco-Americans2018. 388 p.

54. Atlas. Atlas Historique du Canada. La transformation du territoire 1800 - 1891. In: Gentilcore RE., and Robert, J. C.,, editor. Montreal: Les Presses de l’Université de Montréal; 1993. p. 1986.

55. Frenette Y, Rivard É, St-Hilaire M. La francophonie nord-américaine. Québec: Presses de l’Université Laval; 2012.

56. Gagnon A, Miller MS, Hallman SA, Bourbeau R, Herring DA, Earn DJ, et al. Age-specific mortality during the 1918 influenza pandemic: unravelling the mystery of high young adult mortality. PLoS One. 2013;8(8):e69586.

57. Havard G, Vidal, C. Histoire de l’Amérique française: Flammarion; 2014. 863 p.

58. Carvajal-Carmona LG, Soto ID, Pineda N, Ortiz-Barrientos D, Duque C, Ospina-Duque J, et al. Strong Amerind/white sex bias and a possible Sephardic contribution among the founders of a population in northwest Colombia. Am J Hum Genet. 2000;67(5):1287–95.

59. Carvalho-Silva DR, Santos FR, Rocha J, Pena SD. The phylogeography of Brazilian Y-chromosome lineages. Am J Hum Genet. 2001;68(1):281–6.

60. Bedoya G, Montoya P, Garcia J, Soto I, Bourgeois S, Carvajal L, et al. Admixture dynamics in Hispanics: a shift in the nuclear genetic ancestry of a South American population isolate. Proc Natl Acad Sci U S A. 2006;103(19):7234–9.

61. Mendizabal I, Sandoval K, Berniell-Lee G, Calafell F, Salas A, Martinez-Fuentes A, et al. Genetic origin, admixture, and asymmetry in maternal and paternal human lineages in Cuba. BMC Evol Biol. 2008;8:213.

62. Vézina H, Jomphe M, Lavoie E-M, Moreau C, Labuda D. L’apport des données génétiques à la mesure généalogique des origines amérindiennes des Canadiens français. Cahiers québecois de démographie. 2012;41(1):87 – 105.

63. Labuda D, Zietkiewicz E, Labuda M. The genetic clock and the age of the founder effect in growing populations: a lesson from French Canadians and Ashkenazim. Am J Hum Genet. 1997;61(3):768–71.

64. Alves I, Sramkova Hanulova A, Foll M, Excoffier L. Genomic data reveal a complex making of humans. PLoS Genet. 2012;8(7):e1002837.

65. Bouchard G, DeBraekeleer M, editors. Histoire d’un génome. Population et génétique dans l’est du Québec. Québec: Presses de l’Université du Québec; 1991.

66. Scriver CR. Human Genetics: Lessons from Quebec Populations. Annual Review of Genomics and Human Genetics. 2001;2(1):69–101.

67. Laberge AM, Michaud J, Richter A, Lemyre E, Lambert M, Brais B, et al. Population history and its impact on medical genetics in Quebec. Clin Genet. 2005;68(4):287–301.

68. Yotova V, Labuda D, Zietkiewicz E, Gehl D, Lovell A, Lefebvre JF, et al. Anatomy of a founder effect: myotonic dystrophy in Northeastern Quebec. Hum Genet. 2005;117(2-3):177-87.

69. Provine WB. Ernst Mayr: Genetics and speciation. Genetics. 2004;167(3):1041–6.

70. Kere J. Human population genetics: lessons from Finland. Annu Rev Genomics Hum Genet. 2001;2:103–28.

71. Risch N, Tang H, Katzenstein H, Ekstein J. Geographic distribution of disease mutations in the Ashkenazi Jewish population supports genetic drift over selection. Am J Hum Genet. 2003;72(4):812–22.

72. Clegg SM, Degnan SM, Kikkawa J, Moritz C, Estoup A, Owens IP. Genetic consequences of sequential founder events by an island-colonizing bird. Proc Natl Acad Sci U S A. 2002;99(12):8127–32.

73. Tremblay M, Vezina H. Genealogical analysis of maternal and paternal lineages in the Quebec population. Hum Biol. 2010;82(2):179–98.

74. Fix AG. Migration and colonization in human microevolution. Foley RA, Jablonski NG, editors. Cambridge: Cambridge University Press; 1999. 236 p.

75. Reich D. Who we are and how we got here. New York: Pantheon Books; 2018. 335 p.

76. Ramachandran S, Deshpande O, Roseman CC, Rosenberg NA, Feldman MW, Cavalli-Sforza LL. Support from the relationship of genetic and geographic distance in human populations for a serial founder effect originating in Africa. Proc Natl Acad Sci U S A. 2005;102(44):15942–7.

77. Finlayson C. Biogeography and evolution of the genus Homo. Trends Ecol Evol. 2005;20(8):457–63.

78. Cavall-Sforza L, L., Menozzi, Paolo, Piazza, Alberto. The History and Geography of Human Genes. Princeton, NJ: Princeton University Press; 1996.

79. Oppenheimer S. The Real Eve: Modern Man’s Journey Out of Africa. New York: Caroll&Graf Publishers; 2003.

80. Underhill PA, Kivisild T. Use of Y chromosome and mitochondrial DNA population structure in tracing human migrations. Annu Rev Genet. 2007;41:539–64.

81. Alves I, Arenas M, Currat M, Sramkova Hanulova A, Sousa VC, Ray N, et al. Long-Distance Dispersal Shaped Patterns of Human Genetic Diversity in Eurasia. Mol Biol Evol. 2016;33(4):946–58.

82. Lindo J, Haas R, Hofman C, Apata M, Moraga M, Verdugo RA, et al. The genetic prehistory of the Andean highlands 7000 years BP though European contact. Sci Adv. 2018;4(11):eaau4921.

83. Henn BM, Botigue LR, Peischl S, Dupanloup I, Lipatov M, Maples BK, et al. Distance from sub-Saharan Africa predicts mutational load in diverse human genomes. Proc Natl Acad Sci U S A. 2016;113(4):E440–9.

84. Jeong C, Witonsky DB, Basnyat B, Neupane M, Beall CM, Childs G, et al. Detecting past and ongoing natural selection among ethnically Tibetan women at high altitude in Nepal. PLoS Genet. 2018;14(9):e1007650.

85. Peischl S, Dupanloup I, Foucal A, Jomphe M, Bruat V, Grenier JC, et al. Relaxed Selection During a Recent Human Expansion. Genetics. 2018;208(2):763–77.

86. Terwilliger JD, Zollner S, Laan M, Paabo S. Mapping genes through the use of linkage disequilibrium generated by genetic drift: ‘drift mapping’ in small populations with no demographic expansion. Hum Hered. 1998;48(3):138–54.

87. Pouyet F, Aeschbacher S, Thiery A, Excoffier L. Background selection and biased gene conversion affect more than 95% of the human genome and bias demographic inferences. Elife. 2018;7.

88. Martin AR, Gignoux CR, Walters RK, Wojcik GL, Neale BM, Gravel S, et al. Human Demographic History Impacts Genetic Risk Prediction across Diverse Populations. Am J Hum Genet. 2020;107(4):788–9.

89. Gauvreau D, Guerien, M., Hamel, M. De Charlevoix au Saguenay: mesure et caracteristiques du mouvement migratoire avant 1911. In: Bouchard G, De Braekeleer, M., editor. Histoire d’un génome. Québec: Presses de l’Université du Québec; 1991. p. 145–62.

90. Labuda M, Labuda D, Korab-Laskowska M, Cole DE, Zietkiewicz E, Weissenbach J, et al. Linkage disequilibrium analysis in young populations: pseudo-vitamin D-deficiency rickets and the founder effect in French Canadians. Am J Hum Genet. 1996;59(3):633–43.

## References

Larin, R. (2000). Brève histoire du peuplement européen en Nouvelle-France. Sillery (Québec), Septentrion.

Matzuk, M. M. and D. J. Lamb (2002). “Genetic dissection of mammalian fertility pathways.” Nat Cell Biol 4 Suppl: s41–49.

Moreau, C., H. Vezina, V. Yotova, R. Hamon, P. de Knijff, D. Sinnett and D. Labuda (2009). “Genetic heterogeneity in regional populations of Quebec--parental lineages in the Gaspe Peninsula.” Am J Phys Anthropol 139(4): 512–522.

